# Proteomic reprogramming underlies climate-associated variation in seed dormancy and germination of European beech

**DOI:** 10.64898/2026.07.07.736924

**Authors:** Tomasz A. Pawłowski, Marlène Davanture, Andżelika Drozda, Jan Suszka, Mélisande Blein-Nicolas

**Author notes:** Corresponding Author Tomasz A. Pawłowski. Marlène Davanture, Andżelika Drozda, Jan Suszka, Mélisande Blein-Nicolas.

## Abstract

The ability of seeds to survive until dormancy recedes and the germination requirements are met is an adaptive strategy. Proteomics improves our understanding of the mechanisms that control the adaptation to environmental heterogeneity. In this study, we investigated two European beech populations from different habitats that differed in dormancy and germination traits. We found that the populations exhibited different germination strategies, which were reflected in coordinated but quantitatively different proteomic reprogramming. The Miękinia population exhibited stronger accumulation of proteins involved in nucleotide sugar biosynthesis, S-adenosylmethionine metabolism, and flavonoid biosynthesis. Enhanced nucleotide sugar biosynthesis indicates more intensive cell wall remodelling and carbohydrate metabolism, which support embryo growth and faster germination. Increased S-adenosylmethionine metabolism suggests the epigenetic and hormonal regulation of germination differences between populations. Higher flavonoid biosynthesis indicates an enhanced antioxidant capacity associated with environmental protection. In contrast, the Wisła population showed stronger accumulation of proteins involved in RNA processing, suggesting tighter post-transcriptional regulation and proteome reorganization during germination. Consistent with its deeper dormancy and later germination, the Wisła population appears to rely more on RNA-level regulation, whereas the Miękinia population prioritizes metabolic activation. These contrasting proteomic profiles likely reflect population-specific physiological strategies associated with dormancy depth and adaptation to different climatic conditions.

**Highlight:** Proteomic reprogramming reveals population-specific germination strategies in European beech, linking dormancy depth with contrasting metabolic activation and RNA-level regulation during the transition from dormancy to germination.

## Introduction

Over the last few decades, environmental disturbances have affected many aspects of plant life, altering their adaptation strategies and posing challenges to resistance in local habitats (Dyderski *et al*., 2018). According to the regeneration niche theory, the earlier stages of plant development, such as seed dormancy breaking and germination, are the most vulnerable to environmental influences (Rosbakh *et al*., 2018). Researching all aspects of tree regeneration from an ecological perspective is crucial; however, the mechanisms underlying plants’ adaptation potential remain understudied (Kurpisz and Pawłowski, 2022).

Environmental conditions are critical to plant reproduction and control seed dormancy and germination (Penfield, 2017; Klupczyńska and Pawłowski, 2021). Dormancy is an adaptation that coordinates seed germination and plant establishment with the environment. Variation in these seed traits depends on the past climate which selects for locally adapted plant populations. In ecological and evolutionary terms, this property is of paramount importance and is indispensable for preserving species continuity and maintaining biodiversity (Pawłowski *et al*., 2024). Seed dormancy and germination are among the earliest features expressed in the plant life cycle and can critically determine the colonization and distribution of a species (Leuschner and Ellenberg, 2017; Pawłowski *et al*., 2024). The regulation of dormancy and germination initiation enables plants to survive in adverse environmental conditions.

Seed dormancy and germination are tightly regulated developmental processes that determine the timing of seedling establishment, and consequently, the plant’s fitness and the survival success (Sajeev *et al*., 2024). Dormancy is a reversible block to germination that prevents radicle protrusion, even when conditions are otherwise favorable (Hilhorst, 2007). Dormancy release, through processes such as after-ripening, cold stratification, or chemical cues, prepares the embryo to resume the metabolic and developmental programs necessary for germination and seedling growth. Thus, understanding the molecular events underlying the transition from a metabolically quiescent state to a metabolically active and proliferative state is both a fundamental biological issue and a practical necessity for improving plants’ ability to withstand environmental challenges (Tognacca and Botto, 2021).

Proteomics has emerged as a powerful approach for characterizing molecular events in seeds during the transition from dormancy to germination (Baudouin *et al*., 2022; Al-Obaidi *et al*., 2024; Ren and Lv, 2024). This complements transcriptome, epigenome and metabolome studies by providing direct information on the biochemical function effectors, their abundance, processing, and post-translational modifications (Bai *et al*., 2021; Sano *et al*., 2022; Auroux *et al*., 2025). Early comparative proteomic studies in Arabidopsis and other species, including forest species, revealed changes in the proteins abundance associated with carbohydrate and energy metabolism, the translational machinery, gene expression, and signaling during the transition from dormancy to germination (Xia *et al*., 2018; Pawłowski *et al*., 2020; Sghaier-Hammami *et al*., 2020; Pan *et al*., 2023; Pawłowski and Suszka, 2025). These findings suggest that proteome remodeling and selective protein accumulation are key to dormancy alleviation and germination competence.

European beech (*Fagus sylvatica* L.) is a key species in European forests (Bohn *et al*., 2000). Different beech ecotypes have evolved in response to local climatic conditions (Bolte *et al*., 2007; Leuschner and Ellenberg, 2017; Pawłowski *et al*., 2024). This adaptation is also reflected at the seed level (Pawłowski *et al*., 2024). Populations originating from warmer, drier sites have developed deeper dormancy, yet lower germination capacity. However, seed dormancy is surprisingly less influenced by mean annual temperature or precipitation and is more influenced by the length of the frost period, extreme minimum temperature, and evaporation (Pawłowski *et al*., 2024). In this study, we examined the physiological processes of dormancy breaking and germination in two beech populations originating from natural stands in Poland that are characterized by different climatic conditions generally associated with different temperature, precipitation and frost variables (Pawłowski *et al*., 2024). We used quantitative mass spectrometry to systematically profile the proteome dynamics during seed dormancy release and germination in the beech populations. Our aims were threefold: (i) mapping the temporal changes in protein abundance that accompany commitment to germination; (ii) identifying proteins whose regulation correlates with the dormancy status associated with the climate of population’s origin; and (iii) integrating the proteomic signatures with the germination and the population in order to nominate candidate regulators for functional follow-up.

## Materials and Methods

### Beech seed origin and experimental design

In autumn 2020, *Fagus sylvatica* seeds that differed in dormancy depth were collected from two provenances in Poland: Wisła (49°34’19.2“N 18°50’31.0”E, 775 m asl) and Miękinia (50°17’32.0“N 17°27’43.8”E, 380 m asl). Each provenance was represented by five trees. The seeds were dried to a water content of 9% and stored at −3°C. To germinate the seeds, they were imbibed and stratified at 3°C in closed plastic trays and in the dark. Embryo axes were isolated from intact seeds at three time points: dry dormant seeds (D), non-dormant after stratification (ND, without germinated seeds), and germinated (G). For each of the thirty combinations of population, physiological stage, and tree, pools of 50 embryo axes were flash-frozen in liquid nitrogen and stored at −80 °C until further analysis. Prior to the experiment, a germination test was performed on four samples of 50 seeds for each tree according to the ISTA protocol (ISTA, 2019).

### Protein extraction and in solution digestion

The frozen embryo axes were ground into a fine powder using an automatic Tissue Lyser (Qiagen, Venlo, the Netherlands). Proteins were extracted using the phenol extraction protocol described by Faurobert et al. (2007). The dry pellets were then stored at −80 °C prior to being delivered to the PAPPSO proteomics facility. The proteins in the pellets were solubilized in 200 µL of solubilization buffer [6 M urea, 2 M thiourea,10 mM DTT, 30 mM Tris-HCl (pH 8.8), and 0.1% RapiGest SF (Waters)]. Protein concentrations were measured using a 2-D Quant kit (Cytiva). Digestion was performed in 0.2 mL strip tubes. Each sample (40 µg) was alkylated by incubation in the dark for 1 h at room temperature with 2 μL of 300 mM iodoacetamide in 50 mM ammonium bicarbonate. The proteins were then diluted tenfold in 50 mM ammonium bicarbonate and digested using 800 ng of trypsin (enzyme-to-substrate ratio of 1:50 w/w). Digestion was stopped by the addition of 4 μL of 30 % trifluoroacetic acid. The peptides were then desalted using C18 solid-phase extraction (SPE) cartridges (Strata XL, 100 µm, ref. 8E-S043-TGB, Phenomenex, Torrance, CA, USA).

### Mass spectrometry analyses

High-performance liquid chromatography (HPLC) was performed on a nano-HPLC (Vanquish Neo, Thermo Scientific). Buffers A and B were prepared with 0.1% formic acid in water and acetonitrile, respectively. A 4 µL sample of the peptide solution was loaded at a flow rate of 10 µL/min onto an Acclaim PepMap C18 trap column (particle size: 5 µm, pore size: 100 Å, length: 2 cm) and desalted using a solution of 0.1% formic acid and 2% acetonitrile in water. The peptides were then separated on an Acclaim PepMap C18 column (particle size: 2 µm, pore size: 100 Å, length: 25 cm). Peptide separation was achieved at a flow rate of 300 nL/min with the following steps: a linear gradient from 2% to 30% buffer B over 110 min; a linear gradient from 30% to 90% buffer B over 2 min; and regeneration with 90% buffer B over 10 min. The eluted peptides were analyzed online with a Q-Exactive Plus mass spectrometer (Thermo Fisher Scientific, Courtaboeuf, France) using a nanoelectrospray interface. Ionization (with an ionization potential of 1.5 kV) was performed using an uncoated capillary silica tip (360/20-10, New Objective). Peptide ions were analyzed using Xcalibur 4.7.69 with the following data-dependent acquisition steps: (1) a full MS scan (m/z 350–1400 in profile mode) with a resolution of 70,000, and (2) MS/MS (isolation window = 1.5 m/z, AGC target = 1E5, collision energy = 27%, profile mode, resolution = 17,500). Step 2 was repeated for the eight major ions detected in step 1. Dynamic exclusion was set to 50 s. Only doubly, triply, and quadruply charged precursor ions were subjected to MS/MS fragmentation.

### Peptide identification and quantification

The Xcalibur raw data were transformed into the open-source mzML format and centered using the ThermoRawFileParser software (version 1.4.3). Protein identification was performed using X!Tandem PileDriver (version 2015.04.01.1; Craig & Beavis, 2004), which queried the MS/MS data against the UniProt *Fagus sylvatica* protein library (taxonomy ID: 28930), as well as a custom contaminant database (trypsin and keratins). The following static and possible modifications were accounted for: one missed trypsin cleavage, alkylation of cysteine, and oxidation of methionine. Precursor mass tolerance was set to 10 ppm and fragment ion mass tolerance to 0.02 Da. A refinement search was performed with similar parameters, except that possible N-terminal acetylation with peptide signal cleavage was included. Peptide identification and protein inference were performed using i2MassChroQ 1.0.7 (Langella *et al*., 2024) in combine mode. The following criteria were applied: (1) a minimum of two different peptides were required, with an E-value smaller than 0.01; and (2) the protein E-value (calculated as the product of the unique peptide E-values) was smaller than 10⁻⁵. Using a reverse version of the UniProt *Fagus sylvatica* database as a decoy, X!Tandem estimated the false discovery rate (FDR) to be 0.13% and 0.16% for peptide-spectrum matching and peptide identification, respectively. At the protein level, the FDR was estimated to be 0%. Peptide quantification was performed based on extracted ion current (XIC) using MassChroQ (version 2.6; Valot et al., 2011) with the parameters described in Supplementary File S1.

### Data processing and analysis

The peptide quantification data were processed using the MCQR package (version 1.0.2; Balliau et al., 2025), for which an R script is provided in Supplementary File S2. Briefly, the data were filtered to remove seed storage proteins and peptide ions with a retention time standard deviation greater than 20 s or a peak width greater than 100 s. Intensity normalization was performed using the ‘median.RT’ method before filtering for shared peptides, peptide ions showing more than 10% of missing values or showing a correlation coefficient less than 0,5 with the other peptide ions from the same protein. Missing intensity values were imputed using the ‘irmi’ method. Protein abundances were computed from the sum of peptide intensities. Missing protein abundance were imputed. Descriptive analysis of the protein abundance data were performed using principal component analysis and heatmap representation. Hypothesis testing was performed using a two way analysis of variance (ANOVA) followed by a contrast analysis. Functional annotations (Gene Ontology, GO) were obtained from the UniProt for the identified beech proteins. The proteins were then assigned to functional categories using the R software (R Core Team, 2024). This analysis used the ggplot2 (Wickham, 2016), dplyr (Wickham *et al*., 2023), tidyr (Wickham et al., 2024) and stringr (Wickham, 2023) libraries.

The protein sequences were submitted to STRING for homology-based mapping. *Juglans nigra* (*Fagales*) was selected as the closest species with available interaction data. A protein-protein interaction network was constructed using a confidence score of at least 0.4.

## Results

### The Miękinia and Wisła populations have evolved different seed germination parameters

In this study, we examined two beech populations of different origins: Miękinia and Wisła. The Miękinia population grows at lower elevation and is characterized by a shorter frost period and a higher extreme minimum temperature than the Wisła population (Supplementary Table S1). First, we conducted a germination test under controlled conditions to evaluate the differentiation of seed traits between the two populations. We observed that the Miękinia population seeds started to germinate after three weeks, with an average T_50_ of six weeks and a germination capacity of 78%. In contrast, the Wisła population seeds took seven weeks to germinate, with an average T_50_ of ten weeks and a germination capacity of 81% (Figure 1). These results indicate that the Wisła population exhibits deeper dormancy and a higher germination capacity than the Miękinia population, likely due to climatic influences on seed properties.

**Figure 1.**
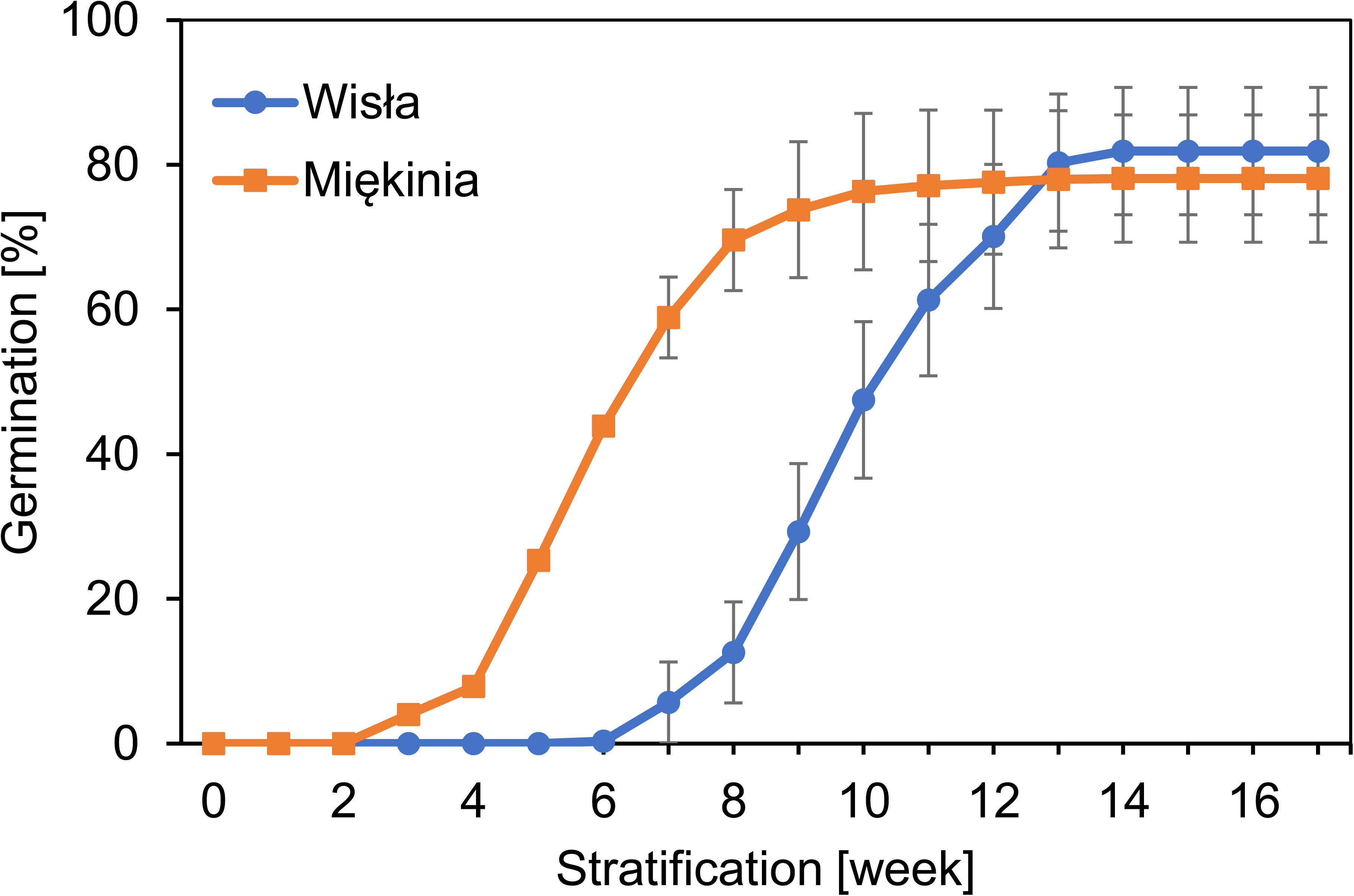
Germination of the beech seeds originating from the Wisła and Miękinia populations after seed imbibition and stratification at 3 °C. Error bars represent standard errors (n = 4; 50 seeds per sample).

### The seed proteome varies depending on the germination stage and the population

To better understand the mechanistic events that drive the environmentally regulated transition from dormancy to active growth in the Miękinia and Wisła beech populations, we conducted a proteomic analysis of the seeds at three germination stages: dry dormant (D), non-dormant after stratification (ND), and germinated (G). A total of 2,916 proteins were quantified. Their abundances in each of the 30 samples are shown in Supplementary Figure S1. As anticipated, given that the seeds were collected in their natural environment, this shows variation in the proteome between trees from the same population and germination stage. There is also variation among germination stages and populations. Principal component analysis (PCA, Figure 2) indicates that germination is the main source of proteome variation (axis 1 explains 52.9% of the total variance). Axis 3 (8.6% of the total variance) primarily separates the two populations, and axis 2 (28.1% of the total variance) mainly distinguishes between Miękinia-D from Wisła-ND. These results suggest the presence of germination-population interactions, as reflected in the first PCA plot, which shows that the two populations are closer at the G stage than the D stage.

**Figure 2.**
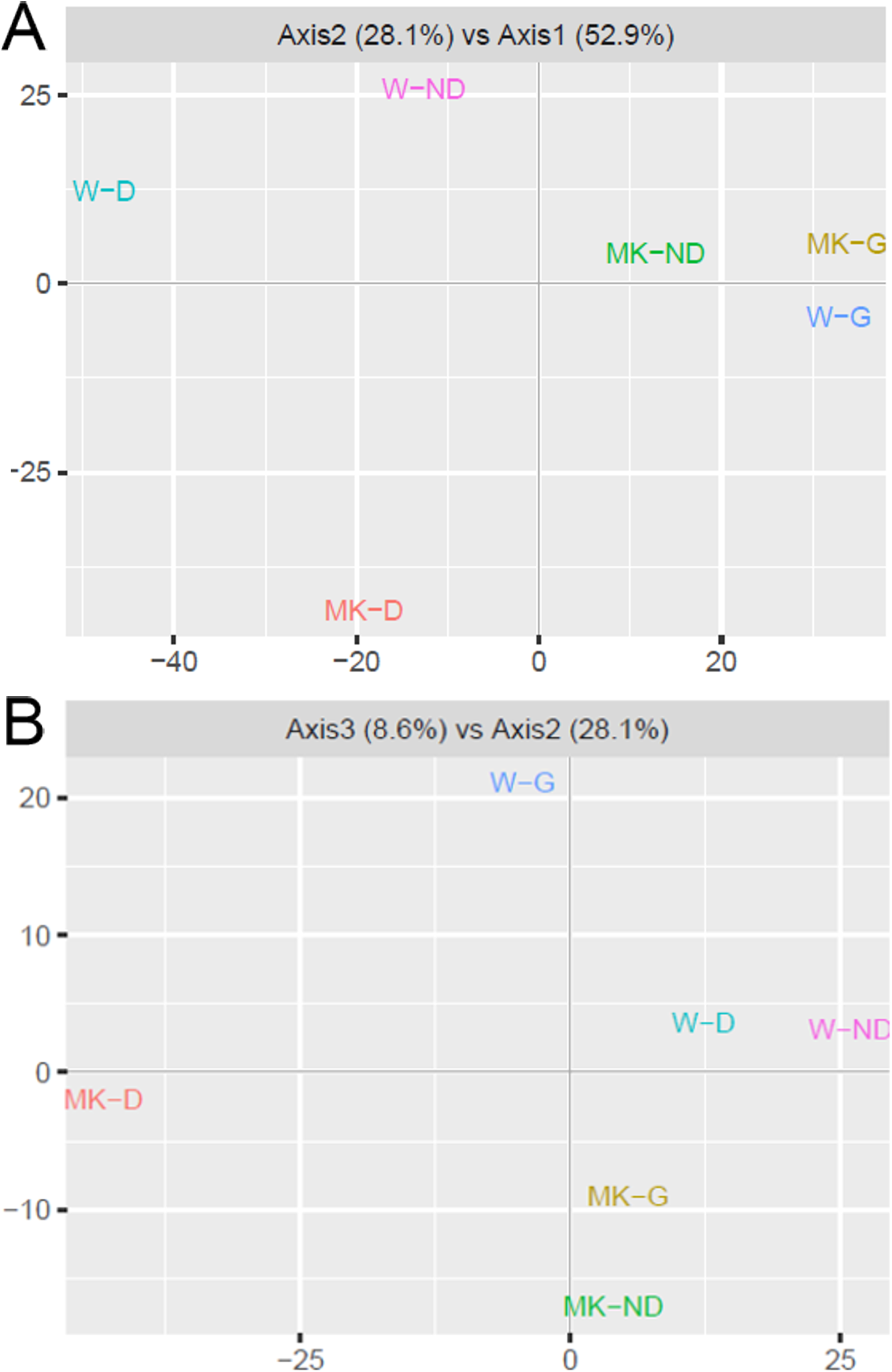
Principal component analysis (PCA) of the effect of germination stages (A) and tree population origins (B) on the proteome of beech seeds. MK – Miękinia, W – Wisła. D - dry dormant seeds, ND - non-dormant seeds after cold stratification, G - germinated seeds.

Next, we searched for proteins showing significant abundance variation using a two-way ANOVA that included germination and population factors, as well as an interaction term (adjusted p-value <0.05). Consistent with the PCA results, we found that germination and population significantly affected the abundance of 1,252 (42.9%) and 450 (15.4%) proteins, respectively (Figure 3). These two sets of proteins overlapped significantly: 79.8% of the proteins that showed a population effect also varied in response to seed germination. However, only 20 proteins exhibited a significant germination x population interaction effect, likely due to the substantial variability between trees and the associated lack of statistical power.

**Figure 3.**
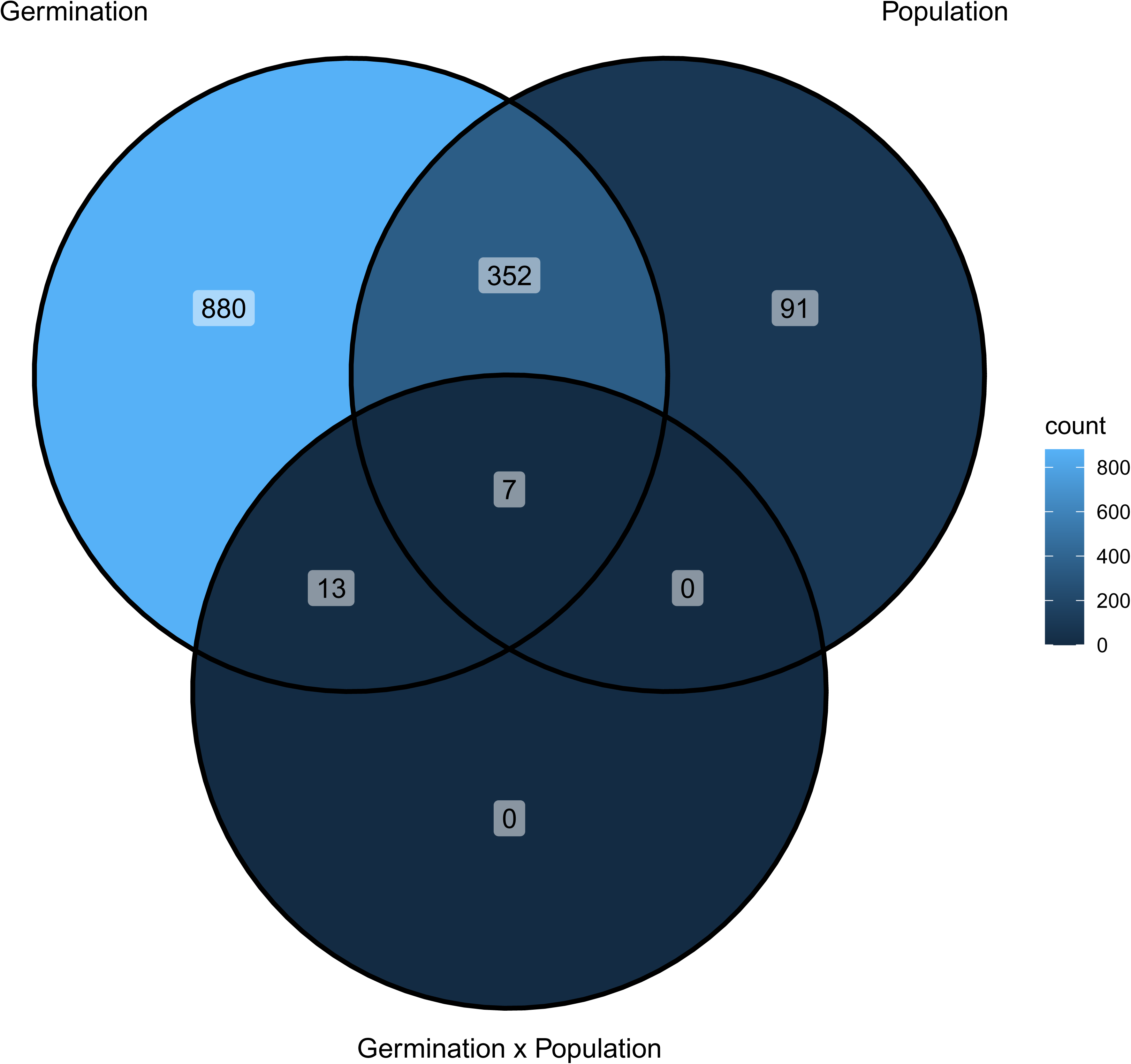
A Venn diagram showing significantly variable beech seed proteins (ANOVA and Tukey test, p < 0.05) for germination and population factors and their interaction.

### Functional analysis of proteins associated with seed dormancy and germination

To better understand the changes in the proteome occurring during seed germination, we analyzed the GO annotations of the 1,252 proteins that responded significantly to the germination stage factor. Besides the unannotated proteins, the majority are associated with the following biological processes (Figure 4): translation (59) proteolysis (37), carbohydrate metabolic processes (27), mRNA processing (19), and protein folding (18). Cellular components include: cytoplasm (153), cytosol (127), nucleus (104), membrane (72) and mitochondria (56). Molecular functions include: ATP binding (136), metal ion binding (93), RNA binding (82), structural constituents of ribosomes (76) and mRNA binding (46).

**Figure 4.**
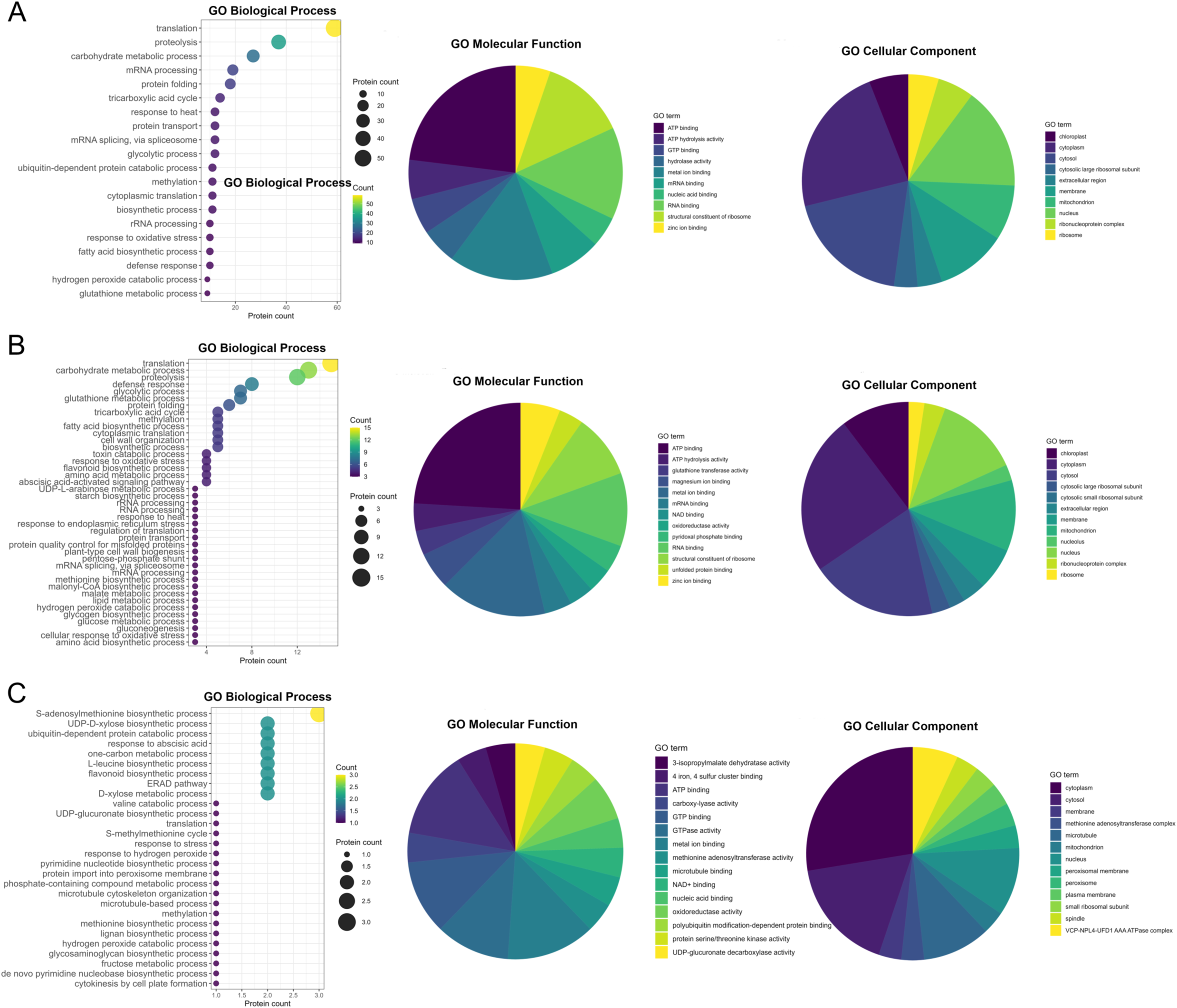
Analysis of Gene Ontology (GO) terms for proteins showing statistically significant changes during beech seed germination (A), between populations (B), and in the interaction between germination and population (C). GO terms are grouped by biological processes, molecular functions, and cellular components.

Then, to gain better insights into the proteome-level transitions occurring between the germination stages, we performed a contrast analysis. This allowed to identify 233 proteins specifically affected by the D to ND transition, 506 proteins specifically affected by the ND to G transition, and 373 proteins affected by both the D to ND and ND to G transitions (Figure 5).

**Figure 5.**
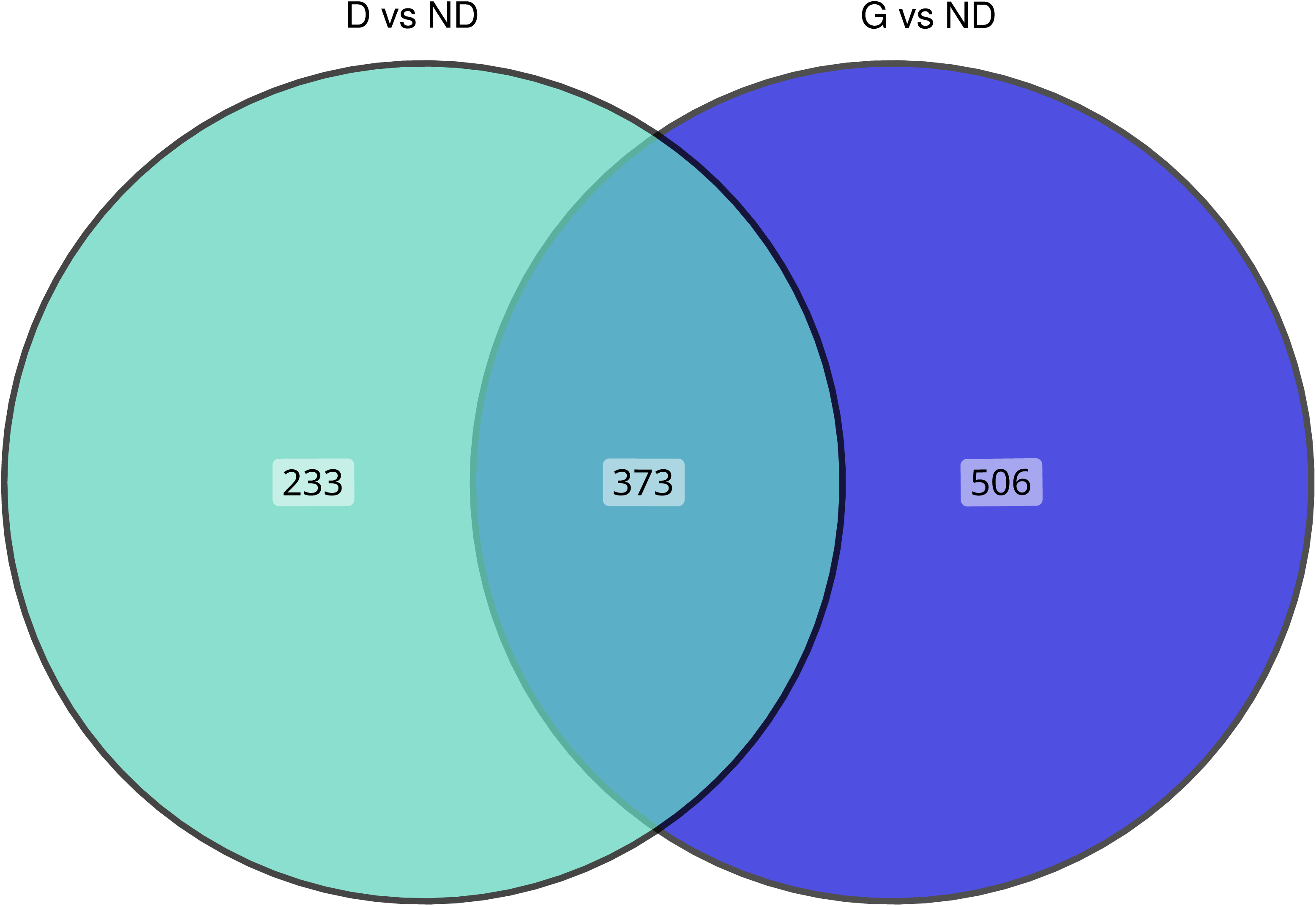
A Venn diagram showing significantly variable beech seed proteins (ANOVA and Tukey test, p < 0.05) across germination stages. D - dry dormant seeds, ND - non-dormant seeds after cold stratification, G - germinated seeds.

### Functional analysis of the proteins associated with seed origin population

A total of 450 proteins exhibited significant abundance variation between the two populations. These proteins were primarily associated with the following GO annotations (Figure 4): biological processes: translation (15), carbohydrate metabolic process (13), proteolysis (12), defense response (8), glutathione metabolic process (7) and glycolytic process (7). Cellular components include: cytoplasm (61); cytosol (48); nucleus (32); mitochondria (27); and chloroplasts (26). Molecular functions include: ATP binding (52), metal ion binding (35), RNA binding (24), structural constituent of ribosome (21) and zinc ion binding (13).

To better understand the proteomic changes that differentiate the two populations during the seed germination process, we performed a contrast analysis focusing on the proteins with significant abundance variation between the two populations at each physiological stage. The results revealed that the proteomes of the Miękinia and Wisła populations differed most significantly at the D stage, with 411 proteins showing significant variation versus 156 and 17 proteins at the ND and G stages, respectively (Figure 6).

**Figure 6.**
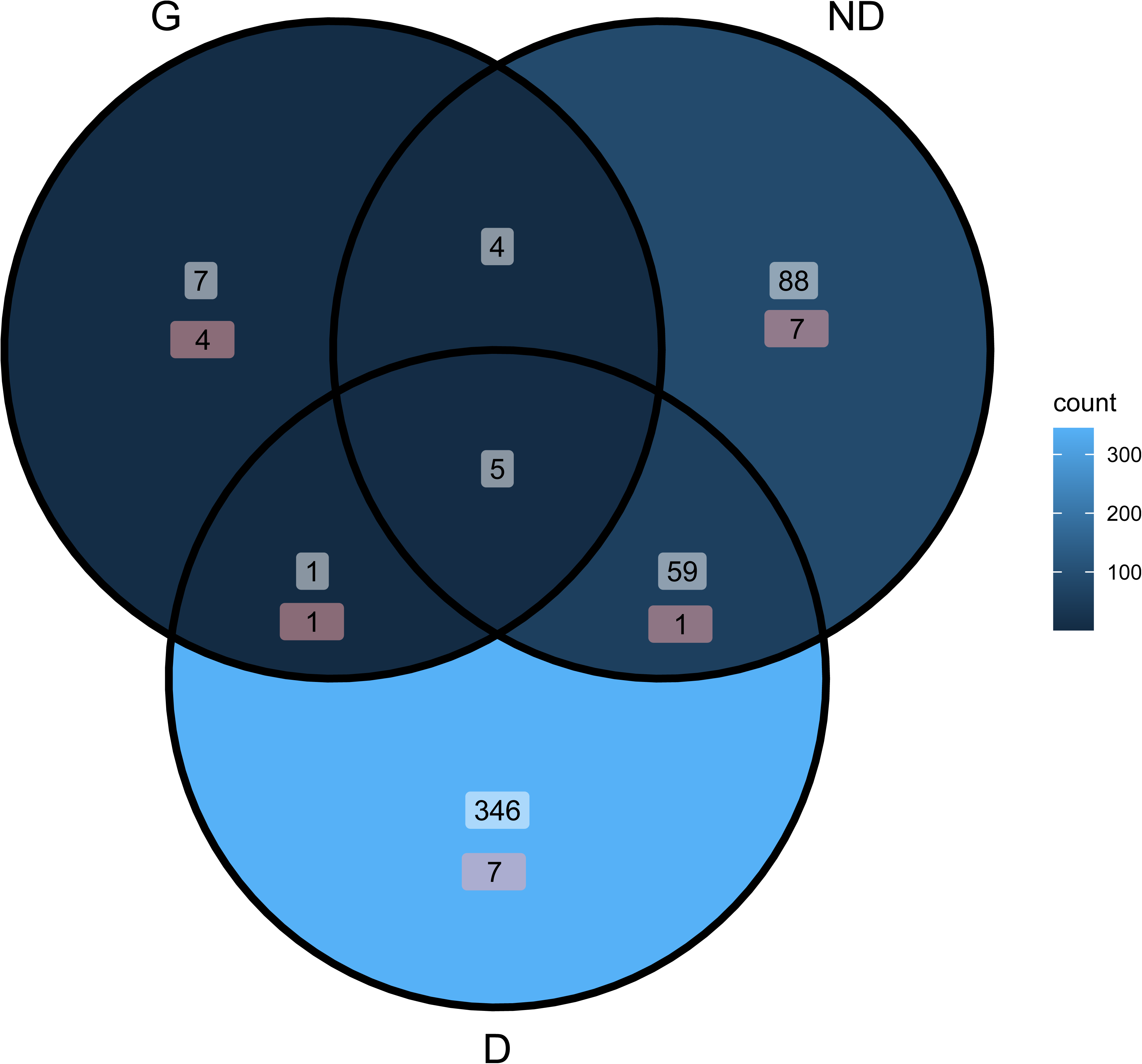
A Venn diagram showing the results of a contrast analysis of the two Miękinia and Wisła populations at dormant, non-dormant, and germinated stages. D - dry dormant seeds, ND - non-dormant seeds after cold stratification, G - germinated seeds.

Of the 20 proteins showing a significant population × stage interaction, seven belonged to the set of 346 proteins that distinguished the Miękinia and Wisła populations specifically at the D stage. Seven more belonged to the group of 88 proteins that differentiated the two populations at the ND stage. Four were included in the set of seven proteins that distinguished the populations at the G stage (Figure 6).

### Functional analysis of the proteins associated with germination and population interaction

To further analyze the interaction between the germination stage and population, we performed a contrast analysis focusing on the 42 proteins for which the abundance difference between populations varied significantly depending on the stage. Twenty-one of these proteins showed a difference in abundance between Miękinia and Wisła that varied between the D and ND stages. Two proteins showed a difference in abundance between Miękinia and Wisła that varied between the ND and D stages. The remaining 19 proteins showed a difference in abundance between Miękinia and Wisła that varied between the D and G stages. Twenty of these proteins were previously identified as significant for the population × germination interaction using ANOVA. Overall, these results suggest that the germination stage x population interaction effects may reflect differences in the physiological state of dormant seeds between the two populations, a disparity that gradually diminishes with stratification and germination.

GO enrichment analysis (Figure 4) of the germination and population interaction contrast analysis showed that the most proteins are associated with the following biological processes: S-adenosylmethionine biosynthetic process (3), D-xylose metabolic process (2), ERAD pathway (2), L-leucine biosynthetic process (2), UDP-D-xylose biosynthetic process (2), flavonoid biosynthetic process (2), one-carbon metabolic process (2), response to abscisic acid (2) and ubiquitin-dependent protein catabolic process (2). Cellular components include cytoplasm (8), cytosol (5), microtubule (3), nucleus (3) and VCP-NPL4-UFD1 AAA ATPase complex (2). Molecular functions include: ATP binding (6), metal ion binding (6), GTP binding (5), GTPase activity (5) and oxidoreductase activity (3).

To further investigate this hypothesis further, we analyzed the protein-protein interaction network of the 42 proteins that showed significant abundance variation between the two populations at each physiological stage. This analysis revealed four functionally coherent modules corresponding to nucleotide-sugar biosynthesis, S-adenosylmethionine (SAM) metabolism, RNA processing and flavonoid biosynthesis (Figure 7). Despite its relatively sparse topology (30 nodes and 14 edges), the network exhibited significant enrichment (p = 0.0108), suggesting nonrandom functional associations among differentially abundant proteins. While all four functional clusters exhibited increased protein accumulation during germination, the relative intensity of this accumulation differed between populations. The Miękinia population showed stronger accumulation of proteins associated with nucleotide-sugar biosynthesis, SAM metabolism, and flavonoid biosynthesis. Conversely, the Wisła population exhibited a more pronounced increase in proteins involved in RNA processing (Figure 8). These results suggest that the two populations employ different germination strategies.

**Figure 7.**
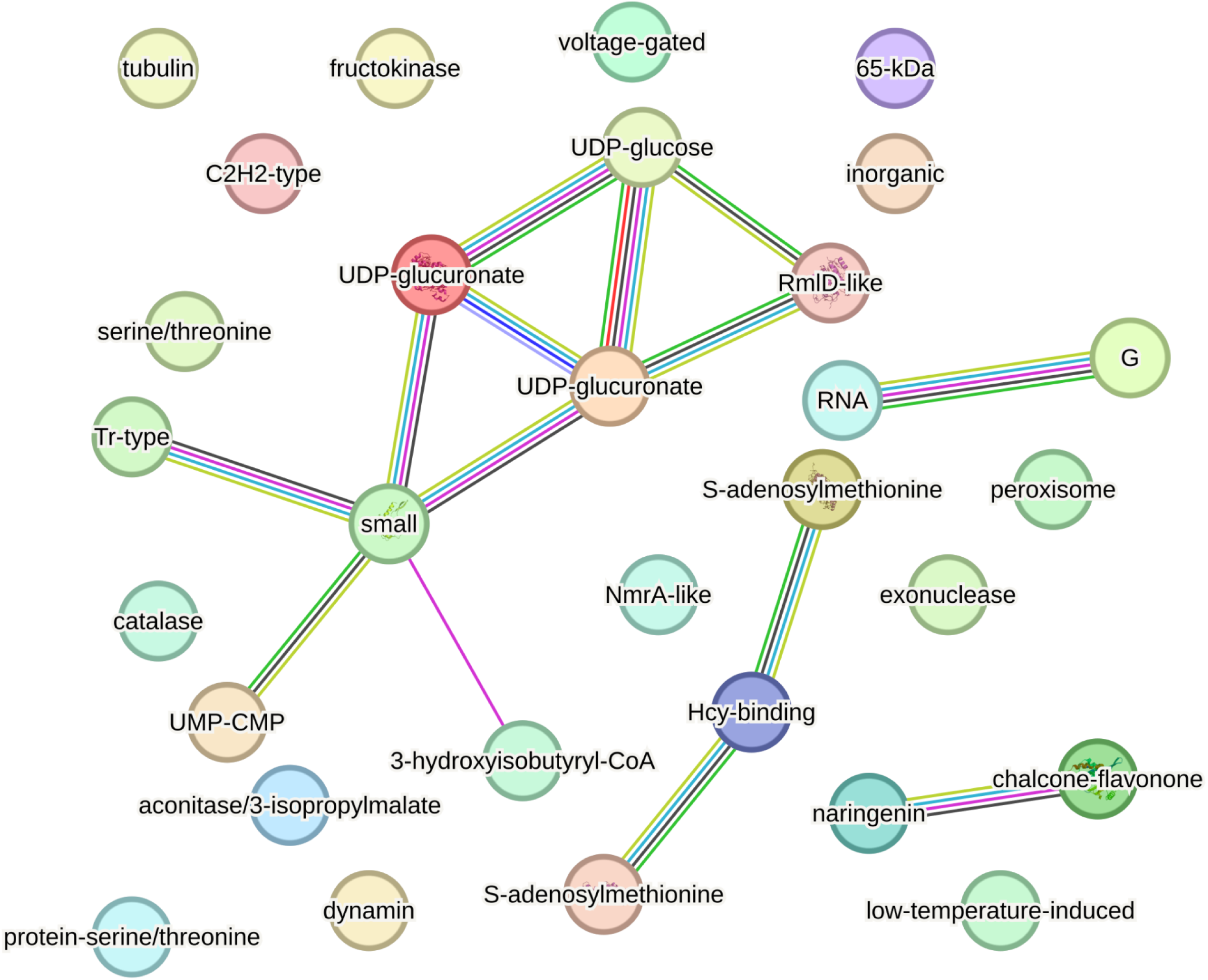
A network diagram showing the protein-protein interactions for beech seed proteins that were found to be differentially abundant after a contrast analysis of the combined germination stages and the two populations.

**Figure 8.**
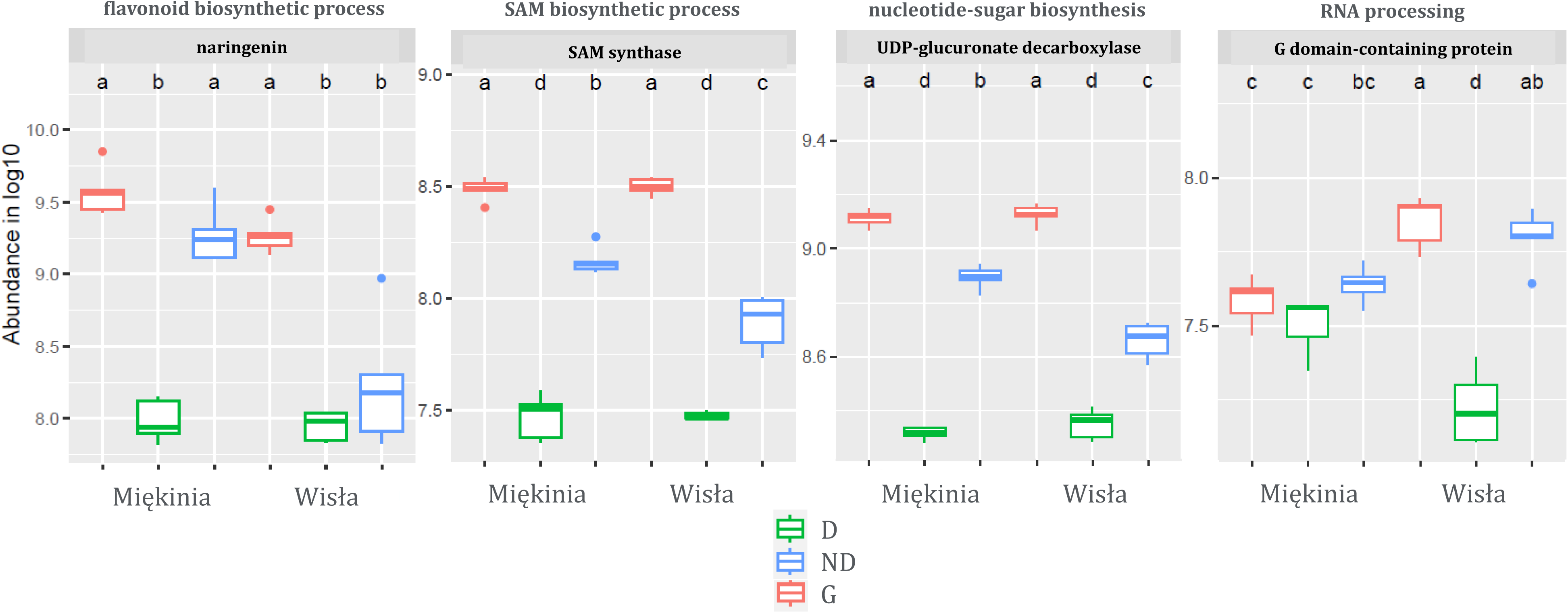
Protein abundance for four functional clusters of beech seed proteins corresponding to nucleotide-sugar biosynthesis, S-adenosylmethionine (SAM) metabolism, RNA processing and flavonoid biosynthesis. D - dry dormant seeds, ND - non-dormant seeds after cold stratification, G - germinated seeds.

## Discussion

Analyzing the proteins involved in the dormancy breaking and germination of beech seeds caused by cold stratification, as well as considering the influence of the beech population’s origin, revealed how trees adapt to the prevailing microclimate. The dormant seed protein pattern differs greatly from the non-dormant and germinated patterns, which are similar to each other. The differences between the dormant and non-dormant stages were greater in the Miękinia population than in the Wisła population. In the Wisła population, the differences between the non-dormant and germinated stages were greater. Dormant seeds from Miękinia population are somewhat similar to non-dormant seeds from the Wisła population, which can be explained by the lower level of initial dormancy in the Miękinia population. These data imply that the populations exhibited highly differential proteome-level expression. As the germination process advanced, the Wisła population exhibited higher protein overexpression than the Miękinia population. Miękinia demonstrated faster preparation for germination than Wisła.

Here, we describe the role of proteins associated with both the germination stage and population origin in regulating beech adaptation mechanism related to germination.

### Nucleotide-sugar biosynthesis associated proteins

The most representative group of proteins was associated with nucleotide-sugar biosynthesis. UDP-glucose 6-dehydrogenase participate in the biosynthesis of UDP-glucuronic acid, while UDP-glucuronate decarboxylase produces UDP-xylose (Reboul *et al*., 2011). This pathway provides nucleotide sugars for the biosynthesis of polysaccharides, including xyloglucans and heteroxylans, which are essential components of the plant cell wall (Harper and Bar-Peled, 2002). Mutations in the UDP-glucuronate decarboxylase gene lead to disturbances in cell wall structure, resulting in abnormalities in seed and leaf development, as well as in vascular tissue conduction (Zhang *et al*., 2005; Litterer *et al*., 2006). UDP-glucose 6-dehydrogenase and UDP-glucuronate decarboxylase activities affect plant growth, defense responses and adaptation to the environment. The present results suggest that these enzymes play a key role in cell wall remodeling during beech seed dormancy breaking and germination, as their levels were found to increase during these processes. The Miękinia population had higher levels of these enzymes than the Wisła population.

UMP-CMP kinase (UMK) catalyzes the phosphorylation of pyrimidine nucleoside monophosphates using ATP as a substrate (Zhou *et al*., 1998). Analysis of Arabidopsis seed germination revealed that the cytosolic UMK3 plays an indispensable role in the biosynthesis of all pyrimidine (deoxy)nucleotides and UDP-sugars (Rinne *et al*., 2024). UMK3 deficiency affects plant growth and development. Mitochondrial UMK2 is essential for dCMP phosphorylation, which is crucial for DNA replication during seed germination (Rinne *et al*., 2024). UMK is also required for the synthesis of coenzymes and energy carriers. The increase in the UMK abundance during the beech seed dormancy breaking and germination demonstrates the pivotal role of nucleotide biosynthesis in this process.

The Tr-type G domain-containing protein is a homolog of the eukaryotic translation initiation factor 5B. Eukaryotic translation initiation factor 5B facilitates rRNA maturation and the transition from translation initiation to elongation, and supports plant development and acclimation to heat stress (Weijers *et al*., 2001; Zhang *et al*., 2017; Wang *et al*., 2019). Other proteins, such as the small ribosomal subunit protein uS7 and the G domain-containing homolog of GTPase (LSG1-2), have been isolated from beech seeds and are involved in ribosome biogenesis and translation during the early stages of plant development (Weis *et al*., 2014). Regulatory crosstalk has been revealed between ribosome biogenesis, mRNA translation initiation, and siRNA biogenesis in plants (Hang *et al*., 2023). Transcription and translation factors, and ribosomal proteins play a crucial role in the gene expression associated with seed dormancy breaking and germination (Pawłowski and Suszka, 2025). The increase in the abundance of these proteins observed in beech seeds during the growth transition and the higher abundance of the G domain-containing protein in the Miękinia population compared to the Wisła population at the dormant stage suggest that gene expression regulation plays a significant role in early plant development and acclimation.

### SAM metabolism associated proteins

SAM biosynthesis proteins have been characterized as being involved in the climate-associated regulation of beech germination. These include SAM synthase, the RmlD-like substrate binding domain-containing protein, and the homocysteine binding domain-containing protein (homocysteine S-methyltransferase) (Roje, 2006; Sauter *et al*., 2013). SAM is essential for DNA and histone methylation, as well as for the biosynthesis of polyamines, and ethylene (Gallardo et al., 2002; Wang et al., 2022). Enzymes involved in SAM biosynthesis strongly influence stress tolerance, epigenetic regulation, and developmental processes including the transition from seed dormancy to germination (Gallardo *et al*., 2002; Kim, 2013; Cohen *et al*., 2017; Wang *et al*., 2022*a*; Lee and Kim, 2024). Homocysteine S-methyltransferase mutant (*mmt*) exhibits reduced germination and growth capacity under stressful conditions (Ogawa and Mitsuya, 2012). The abundance of SAM synthase and the RmlD-like substrate binding domain-containing protein increased during beech seed dormancy breaking and germination, in non-dormant seeds it was lower in Wisła than in Miękinia. These results underscore the significance of SAM metabolism in breaking seed dormancy and germination. Additionally, they imply that population differences may be epigenetically regulated (Pawłowski, 2009, 2010; Pawłowski and Staszak, 2016).

### Flavonoid biosynthesis associated proteins

The flavonoid biosynthetic process was represented by the enzymes chalcone-flavonone isomerase and naringenin 2-oxoglutarate 3-dioxygenase (Shirley *et al*., 1992). Flavonoids play a co-evolutionary role in promoting the environmental adaptability of land plants through their environmentally regulated biosynthesis (Kurepa *et al*., 2023). The flavonol quercetin regulates the abscisic acid (ABA) signaling and the response to environmental severity (Brunetti *et al*., 2018, 2019). Additionally, gibberellic acid (GA) promotes root growth by directly reducing flavonol biosynthesis with the participation of DELLA proteins (Tan *et al*., 2019). An increase in these two proteins was observed during beech seed dormancy breaking and germination, with a higher abundance in the Miękinia populations at the non-dormant stage. Our results imply that flavonoid biosynthetic enzymes play a crucial role in adapting plant development to the environmental diversity. This is particularly significant in the Miękinia population, which, according to the presented results, allocates more resources to protective components.

### RNA processing associated proteins

RNA helicases are essential ATP-dependent enzymes that remodel RNA and regulate its processing and translation into proteins (Li *et al*., 2023). In Arabidopsis, RH12 positively regulates seed germination under stress conditions by reducing reactive oxygen species (ROS) accumulation and promoting the decay of germination-related mRNA transcripts, which involves a DEAD-box helicase (Yuan *et al*., 2024). The RNA helicase RH12 also negatively regulates the ABA pathway. The helicase RH58 enhances seed germination under salt or dehydration stress (Nawaz and Kang, 2019). It has been showed that the DEAD-box RNA helicase 22 is necessary for proper accumulation of mRNA of plastid genes during seed development and seedling growth (Kanai *et al*., 2013). The RNA helicase identified in the present study was positively regulated during the release of dormancy and germination of beech seeds. It is thought to be involved in expressing the RNA necessary for establishing seedlings.

### Other function proteins

The voltage-gated potassium channel is a membrane protein that mediates the transport of potassium ions (K^+^) (Tang *et al*., 1996). K^+^ is important for enzyme activation, stabilization of protein synthesis, neutralization of negative charges on proteins, formation of the membrane potential, and maintenance of cytosolic pH homeostasis (Dreyer and Uozumi, 2011; Dreyer *et al*., 2021; Lefoulon, 2021). K^+^ channels are responsible for potassium uptake and transport. Through K^+^ sequestration, they control stomata movement, growth and cell expansion, stress responses and signaling (Schachtman, 2000; Hendrich, 2012; Sharma *et al*., 2013). During Arabidopsis seed germination, the two-pore K (*TPK1*) potassium channel gene contributes to radicle development by depositing K^+^ in the vacuole to promote expansion growth (Gobert *et al*., 2007). Another potassium channel protein, ZmKCH5, positively mediates the rate of seed germination and primary root growth under salt stress (Zhang *et al*., 2025). The increased abundance of this protein in beech seeds between the dormant and non-dormant and germination stages, implies its importance of this protein in K⁺ transport and signaling, as well as its positive impact on embryo growth.

Aconitase/3-isopropylmalate dehydratase belongs to the aconitase superfamily, and has structural and mechanistic similarities to 3-isopropylmalate dehydratase (Imhof *et al*., 2014). In mitochondria, aconitase functions in the tricarboxylic acid cycle. In contrast, cytosolic aconitase contributes to the glyoxylate cycle and glutamate biosynthesis (Hooks *et al*., 2014). Genetic divergence in aconitase allows plants to adjust their metabolism in response to different developmental stages and environmental stimuli, enhancing their adaptability and resilience (Wang *et al*., 2016*b*; Rahikainen *et al*., 2025). Aconitase is essential for seed germination thus providing energy from lipid reserves to the emerging seedling (Stephen *et al*., 2004; Hooks *et al*., 2014; De Bellis *et al*., 2020). 3-Isopropylmalate dehydratase is involved in leucine and methionine biosynthesis (He *et al*., 2011). Plants in which the *SSU1* gene (which codes for 3-isopropylmalate dehydratase) has been silenced exhibit stunted growth, abnormal morphology, and disordered seed development (Lächler *et al*., 2020). The decrease in protein abundance in beech seeds between the dormant and non-dormant stages and the increase at the germination stage, implies the importance of this protein in processes associated with seed maturation and germination, such as reserve mobilization and protein synthesis. Differential abundance between populations, higher abundance in the Wisła population in the dormant state, shows the populations’ differential adaptation to their environments.

The 65-kDa microtubule-associated protein 1 (MAP65-1) and the tubulin beta chain are involved in mitosis and cytokinesis through the organization of the microtubule cytoskeleton (Smertenko *et al*., 2006; Vavrdová *et al*., 2019). They are essential for seed germination and seedling establishment through by participating in cell division, expansion and directional growth (Pawłowski *et al*., 2004; Yan *et al*., 2020; Kim *et al*., 2024). In interphase cells, MAP65-1 and MAP65-2 help form cortical microtubule arrays that guide directional cell expansion (Kim *et al*., 2024). MAP65-1 expression increases during Arabidopsis seed dormancy breaking and germination, especially upon exposure to GA₃ (Kim *et al*., 2024). Additionally, *At*MAP65-1 and *Gm*MAP65-1 confer tolerance of plants to various biotic and abiotic stresses including cold tolerance (Kim *et al*., 2024). An increase in MAP65-1 and tubulin beta chain abundance was observed during dormancy breaking and germination in beech seeds. The Miękinia population had higher levels than the Wisła population at the non-dormant stage, which was associated with a faster preparation for germination in the Miękinia population. In summary, MAP65-1 and beta-tubulin ensure proper cell division and directed cell growth during seed germination, even in adverse environments.

The C2H2-type domain-containing proteins, also known as C2H2 zinc finger proteins (ZFPs), are associated with the endoplasmic reticulum--associated protein degradation (ERAD) pathway and the ubiquitin-dependent protein catabolic process. These proteins regulate gene transcription in plants and can also mediate the ubiquitin-dependent degradation of target proteins by interacting with co-repressors such as TOPLESS. TOPLESS acts as an adapter for E3 ubiquitin ligase complexes (Lyu and Cao, 2018; Han *et al*., 2020). Arabidopsis ZFP3 acts as a negative regulator of ABA signaling during germination (Joseph *et al*., 2014). In rice, OsZFP15 accelerates germination and increases salt and drought tolerance by influencing ABA catabolism (Wang *et al*., 2022*c*). The increase in protein abundance observed during the beech seed dormancy breaking suggests that ZFPs may abolish the inhibitory effect of ABA, thereby causing germination.

Inorganic diphosphatase (pyrophosphatase, PPA) has two functions in plant cells. First, it hydrolyzes cytosolic pyrophosphate to maintain its level. Second, it translocates protons into vacuoles to maintain the acidity of the vacuolar lumen (Ferjani *et al*., 2012; Segami *et al*., 2018). Through that PPA controls the equilibrium of gluconeogenic reactions during the heterotrophic growth phase of early seedling establishment (Meyer *et al*., 2012). PPA also determines the rate of cytosolic glycolysis, providing carbon for lipid accumulation during seed maturation (Meyer *et al*., 2012). PPA plays an important role in plant tolerance to salt and drought stress during Arabidopsis seed germination (Liu *et al*., 2011). The increase in PPA abundance observed during beech seed germination suggests that this enzyme is crucial to the process. Its function can be associated with supplying energy and protecting embryo tissues against stress. Interestingly, PPA was found to be more abundant in the Miękinia than in the Wisła population during the non-dormant and germinated stages.

Dynamin GTPase, also known as dynamin-related protein (DRP) is an enzyme involved in the vesicle trafficking from the trans-Golgi network to the central vacuole (Jin *et al*., 2001). In plants, the function of dynamins is associated with membrane remodeling, organelle biogenesis, endocytosis, cytokinesis and cell expansion (Bednarek and Backues, 2010). An Arabidopsis mutant of the dynamin gene, *drp1AWS*, exhibited early seedling arrest (Collings *et al*., 2008). *Van3* (another component of the vesicle trafficking) and *drp1AWS* double mutants fail to germinate and exhibit enhanced defects in vascular structure (Sawa *et al*., 2005). The cell plate restricted association of DRP1A and auxin carrier PIN proteins has been found to be necessary for establishing cell polarity and further cytokinesis in Arabidopsis (Mravec *et al*., 2011). An increase in dynamin GTPase abundance was observed during beech seed dormancy breaking and germination. These results suggest that auxin-mediated vesicle trafficking plays an important role in seed germination.

Fructokinase channels fructose into glycolysis, the hexose-phosphate pools and the synthesis of UDP-sugars, all of which are necessary for growth (Stein and Granot, 2018). During germination fructokinase helps to convert the liberated fructose from storage reserves into the metabolites required for respiration and biosynthesis to fuel embryo growth. Metabolic profiling of germinating seeds revealed a significant shift toward hexose phosphorylation and glycolysis (Fait *et al*., 2006). Tomato fructokinase mutants had very low germination rates (Lugassi *et al*., 2022). Our results also confirm the important role of fructokinase in seed germination, as its abundance increased when beech seeds broke dormancy and germinated.

The exonuclease domain-containing protein showed homology with the small RNA-degrading nuclease 3 (SDN3). SDN3 contributes to the 3’-5’ exonuclease degradation of microRNAs bound to ARGONAUTE1 (AGO1) (Ramachandran and Chen, 2008; Chen *et al*., 2018). This process regulates RNA silencing pathways. Since miRNAs regulate key transcription factors, SDN activity indirectly affects plant development, hormone signaling, and stress responses (Yu *et al*., 2017). The direct role of SDN3 in seed germination remains unclear. However, since miRNA activity controls seed dormancy, ABA signaling, and early seedling development, SDN3 likely plays a role in shaping the miRNA landscape during the transition from dormancy to germination (Liu *et al*., 2007; Reyes and Chua, 2007; Das *et al*., 2015; Jiang *et al*., 2022). Since the abundance of the beech seed exonuclease increases during dormancy breaking and germination, one could hypothesize that it reduces the level of miRNAs that block the expression of some germination-positive genes.

The serine/threonine protein kinase (STK) catalyzes the transfer of phosphate groups in signal cascades and plays a fundamental role in cell development and auxin signaling (Wingenter *et al*., 2011; Jagodzik *et al*., 2018). STK is a paralog of the ABA-regulated, seed-expressed serine/threonine kinase family found in beech and known as FsPKs (Lorenzo *et al*., 2003; Jiménez *et al*., 2006; Reyes *et al*., 2006). FsPKs function in maintaining seed dormancy (Lorenzo *et al*., 2003; Jiménez *et al*., 2006; Reyes *et al*., 2006). Other plant STKs, SnRK2s, play a central role in regulating germination by balancing the ABA–GA pathways and integrating stress signals (Nakashima *et al*., 2009; Fujii *et al*., 2011). SnRK2s promote plant growth under optimal conditions and in the absence of ABA, but inhibit the growth of plants in response to ABA (Hasan *et al*., 2022). In summary, STKs are the key regulators of seed germination (Wu *et al*., 2025). Serine/threonine protein phosphatase (PP) counterbalances the activity of STKs, modulating ABA and auxin pathways, MAPK, and calcium signaling (Ghanizadeh *et al*., 2025). PP2Cs dephosphorylate and inhibit SnRK2 kinases, thereby regulating the stress response and inhibiting seed germination (Yoshida *et al*., 2006; Ghanizadeh *et al*., 2025). In the case of beech seeds, STKs increased in abundance during the dormancy breaking and germination and were present at higher level in non-dormant and germinated seeds in the Miękinia population. PP had the opposite effect: its level decreased as germination progressed. This study demonstrates the important roles played by STKs and PPs in coordinating plant development and adaptive responses to environmental challenges.

Peroxisome biogenesis proteins, commonly referred to as peroxins (PEXs), are essential for the formation, maintenance, and regulation of peroxisomes in plant cells (Hu, 2007; Hu *et al*., 2012). These dynamic organelles are involved in various metabolic pathways, including the β-oxidation of fatty acids, detoxification of reactive oxygen species, and the biosynthesis of auxin, jasmonate and salicylic acid (Kao *et al*., 2018; Muhammad *et al*., 2022). These pathways are particularly active during seed germination and seedling establishment (Desai and Hu, 2008). PEXs ensure the rapid adaptation of peroxisome activity to developmental and environmental cues (Muhammad *et al*., 2022). The increase in PEX abundance in beech seeds from the dormant stage to germination underscores the importance of peroxisome-associated processes in germination regulation. Differences in abundance between populations at the dormant stage, higher in the Wisła population, demonstrate how climatic conditions at the seeds’ place of origin influence seed physiology.

The low-temperature-induced 65 kDa protein is a cold-responsive protein that typically accumulates during cold acclimation to help plants survive and adapt to chilling or freezing stress (Pang *et al*., 2023; Fang *et al*., 2025). The interaction between the ABA receptor PYL8, ABI5 and the low-temperature-induced 65 kDa protein indicates that the latter plays a role in breaking seed dormancy in *Malus sieversii* (Fang *et al*., 2025). The highest level of this protein was found in beech dormant seeds, suggesting that it protect against low temperatures and is involved in acquiring or maintaining dormancy.

3-Hydroxyisobutyryl-CoA hydrolase (CHY) catalyzes the final steps of the valine catabolic process (Taylor *et al*., 2004). In the mitochondria, CHY converts valine into propionyl-CoA, which is then metabolized into succinyl-CoA, an intermediate in the tricarboxylic acid cycle (Taylor *et al*., 2004). This ultimately links amino acid catabolism to cellular energy and carbon metabolism, particularly in situations of metabolic stress (Gipson *et al*., 2017). Disruptions in the mitochondrial valine degradation pathway have been found to affect Arabidopsis seed development and germination (Gipson *et al*., 2017). CHY4 plays an essential role in embryo development (Gipson *et al*., 2017). The increased level of CHY in beech seeds during dormancy release and germination suggests that it plays a role in both embryo development and growth.

Catalase catalyzes the breakdown of ROS, specifically converting hydrogen peroxide (H_2_O_2_) into water and oxygen (Wojtyla *et al*., 2016). This protects cells from the toxic effects of ROS. As seeds transition from dormancy to germination, increased respiration and metabolic activity result in a surge of ROS (Wojtyla *et al*., 2016). If these ROS are not efficiently removed, they can be harmful in excess. Additionally, ROS can also play a second role in seed physiology by acting as cellular signaling agents that promote dormancy breaking and germination (Bailly *et al*., 2008). Arabidopsis ABI5 regulates ROS homeostasis by activating CATALASE 1 transcription during seed germination (Bi *et al*., 2017). ROS homeostasis plays a role in how seeds perceive environmental factors by seeds during germination and thus controls its completion (Bailly *et al*., 2008). The observed increase in beech seeds during dormancy release and the subsequent decrease during germination suggests that catalase plays a protective or regulatory role in breaking dormancy.

The NmrA-like domain-containing protein is involved in the lignan biosynthesis process and exhibits pinoresinol-lariciresinol reductase (PLR) activity (Nakatsubo *et al*., 2008). Lignans provide structural support and protect against pathogens and abiotic stress (Markulin *et al*., 2019*a*). They also regulate plant growth and lignification processes, particularly in woody plants (Singh *et al*., 2024). Lignan biosynthesis is positively controlled by ABA and negatively by GA, via the transcriptional regulation of the *PLR1* gene (Renouard *et al*., 2012; Corbin *et al*., 2013; Markulin *et al*., 2019*b*). PLR regulates phenylpropanoid metabolism, including the production of lignin, flavonoids and other phenolic compounds (Markulin *et al*., 2019*a*). PLR activity is associated with seed development and increases during germination (Wang *et al*., 2016*a*; Bekele *et al*., 2025). We found that the abundance of the NmrA-like domain-containing protein increased during the release of beech seed dormancy and germination. Its abundance was higher in the Miękinia population than in the Wisła population. Its role is probably related to the protection of early-growing beech tissues.

### Conclusions

Both populations activated the same four functional modules during germination. This is reflected by increased protein accumulation in nucleotide-sugar biosynthesis, SAM metabolism, RNA processing, and flavonoid biosynthesis. However, the relative intensity of these processes differed, indicating distinct germination strategies. The Miękinia population exhibited stronger accumulation of proteins involved in nucleotide-sugar biosynthesis, SAM metabolism, and flavonoid biosynthesis. This pattern suggests a strategy characterized by rapid metabolic activation, enhanced cell wall remodeling, and increased secondary metabolism. This is consistent with the population’s faster germination and shallower dormancy. In contrast, the Wisła population exhibited a stronger increase in proteins related to RNA processing. Given its deeper dormancy and longer stratification requirement, this pattern may indicate tighter post-transcriptional control and more regulated proteome reorganization during the transition from dormancy to active growth. Thus, while the two populations share the core germination machinery, they differ in their relative investment in metabolic versus RNA-level regulation. These results support the presence of population-specific proteomic reprogramming underlying divergent germination dynamics.

## Abbreviations

ABA: abscisic acid
CHY: 3-hydroxyisobutyryl-CoA hydrolase
D: dormant seeds
DRP: dynamin-related protein
G: germinated seeds
GA: gibberellic acid
GO: gene ontology
MAP65-1: 65-kDa microtubule-associated protein 1
N: non-dormant seeds
PLR: pinoresinol-lariciresinol reductase
PP: serine/threonine protein phosphatase
PPA: pyrophosphatase
PEX: peroxin
ROS: reactive oxygen species
SAM: S-adenosylmethionine
SDN3: small RNA-degrading nuclease 3
STK: serine/threonine protein kinase
UMK: UMP-CMP kinase
ZFP: C2H2 zinc finger proteins

## Supplementary data

The following supplementary data are available at JXB online.

**Table S1.** Population data including climatic variables of studied *Fagus sylvatica* populations. The data were derived from ClimateEU (Marchi *et al*., 2020) for the period from 1991 to 2020.

**Figure S1.** A heatmap showing significant changes to the beech seed proteome during germination in two populations, with hierarchical clustering. Mk – Miękinia, W – Wisła. D - dry dormant seeds, ND - non-dormant seeds after cold stratification, G - germinated seeds.

**File S1.** R script for peptide quantification parameters based on extracted ion current (XIC) using MassChroQ (version 2.6; Valot et al., 2011).

**File S2.** R script for processing the peptide quantification data using the MCQR package (version 1.0.2; Balliau et al., 2025).

## Acknowledgements

We are grateful to Barbara Kurpisz, Elżbieta Nogajewska, Magdalena Sobczak, and Danuta Szymańska for collecting and maintaining the materials for the experiment.

## Author contributions

TAP: conceptualization; TAP, MD, MB-N: methodology; TAP, MD, MB-N: formal analysis; TAP, MD, MB-N: investigation; TAP and JS: resources; TAP, MD, MB-N: data curation; TAP: writing - original draft; TAP, MD, AD, JS and MB-N: writing - review & editing; TAP, MD, MB-N: visualization; TAP: supervision; TAP: funding acquisition.

## Data availability

The mass spectra data underlying this article are available in [repository name] at [URL], and can be accessed with [unique identifier, e.g. accession number, deposition number] (in progress)

## References

1. Al-Obaidi JR, Lau S-E, Liew YJM, Tan BC, Rahmad N. 2024. Unravelling the Significance of Seed Proteomics: Insights into Seed Development, Function, and Agricultural Applications. The Protein Journal 43, 1083–1103.

2. Auroux L, Liew LC, Whelan J, Lewsey MG. 2025. Advances in seed omics. Journal of Experimental Botany, eraf294.

3. Bai B, van der Horst N, Cordewener JH, America AHP, Nijveen H, Bentsink L. 2021. Delayed Protein Changes During Seed Germination. Frontiers in Plant Science 12.

4. Bailly C, El-Maarouf-Bouteau H, Corbineau F. 2008. From intracellular signaling networks to cell death: the dual role of reactive oxygen species in seed physiology. Comptes Rendus Biologies 331, 806–814.

5. Balliau T, Frambourg A, Langella O, Martin M-L, Zivy M, Blein-Nicolas M. 2025. MCQR: Enhancing the Processing and Analysis of Quantitative Proteomics Data by Incorporating Chromatography and Mass Spectrometry Information. Journal of Proteome Research 24, 2861–2873.

6. Baudouin E, Puyaubert J, Meimoun P, Blein-Nicolas M, Davanture M, Zivy M, Bailly C. 2022. Dynamics of Protein Phosphorylation during Arabidopsis Seed Germination. International Journal of Molecular Sciences 23, 7059.

7. Bednarek SY, Backues SK. 2010. Plant Dynamin-Related Protein Families DRP1 and DRP2 in Plant Development. Biochemical Society transactions 38, 797–806.

8. Bekele B, Gallach M, Mekonen T, Beyene D, Tesfaye K, Andargie M. 2025. Integrative metabolomic and transcriptomic analyses reveal key mechanisms of lignan biosynthesis during sesame (Sesamum indicum L.) seed development. BMC Plant Biology 25, 1467.

9. Bi C, Ma Y, Wu Z, Yu Y-T, Liang S, Lu K, Wang X-F. 2017. Arabidopsis ABI5 plays a role in regulating ROS homeostasis by activating CATALASE 1 transcription in seed germination. Plant Molecular Biology 94, 197–213.

10. Bohn U, Gollub G, Hettwer C, Weber H, Neuhäuslová Z, Raus T, Schlüter H. 2000. Karte der natürlichen Vegetation Europas / Map of the Natural Vegetation of Europe - Maßstab / Scale 1:2,500,000.

11. Bolte A, Czajkowski T, Kompa T. 2007. The north-eastern distribution range of European beech—a review. Forestry 80, 413–429.

12. Brunetti C, Fini A, Sebastiani F, Gori A, Tattini M. 2018. Modulation of Phytohormone Signaling: A Primary Function of Flavonoids in Plant–Environment Interactions. Frontiers in Plant Science 9.

13. Brunetti C, Sebastiani F, Tattini M. 2019. Review: ABA, flavonols, and the evolvability of land plants. Plant Science 280, 448–454.

14. Chen J, Liu L, You C, et al. 2018. Structural and biochemical insights into small RNA 3t end trimming by Arabidopsis SDN1. Nature Communications 9, 3585.

15. Cohen H, Salmon A, Tietel Z, Hacham Y, Amir R. 2017. The relative contribution of genes operating in the S-methylmethionine cycle to methionine metabolism in Arabidopsis seeds. Plant Cell Reports 36, 731–743.

16. Collings DA, Gebbie LK, Howles PA, Hurley UA, Birch RJ, Cork AH, Hocart CH, Arioli T, Williamson RE. 2008. Arabidopsis dynamin-like protein DRP1A: a null mutant with widespread defects in endocytosis, cellulose synthesis, cytokinesis, and cell expansion. Journal of Experimental Botany 59, 361–376.

17. Corbin C, Renouard S, Lopez T, Lamblin F, Lainé E, Hano C. 2013. Identification and characterization of cis-acting elements involved in the regulation of ABA- and/or GA-mediated LuPLR1 gene expression and lignan biosynthesis in flax (Linum usitatissimum L.) cell cultures. Journal of Plant Physiology 170, 516–522.

18. Craig R, Beavis RC. 2004. TANDEM: matching proteins with tandem mass spectra. Bioinformatics 20, 1466–1467.

19. Das SS, Karmakar P, Nandi AK, Sanan-Mishra N. 2015. Small RNA mediated regulation of seed germination. Frontiers in Plant Science 6, 828.

20. De Bellis L, Luvisi A, Alpi A. 2020. Aconitase: To Be or not to Be Inside Plant Glyoxysomes, That Is the Question. Biology 9.

21. Desai M, Hu J. 2008. Light Induces Peroxisome Proliferation in Arabidopsis Seedlings through the Photoreceptor Phytochrome A, the Transcription Factor HY5 HOMOLOG, and the Peroxisomal Protein PEROXIN11b. Plant Physiology 146, 1117–1127.

22. Dreyer I, Sussmilch FC, Fukushima K, Riadi G, Becker D, Schultz J, Hedrich R. 2021. How to Grow a Tree: Plant Voltage-Dependent Cation Channels in the Spotlight of Evolution. Trends in Plant Science 26, 41–52.

23. Dreyer I, Uozumi N. 2011. Potassium channels in plant cells. The FEBS Journal 278, 4293–4303.

24. Dyderski MK, Paz S, Frelich LE, Jagodzinski AM. 2018. How much does climate change threaten European forest tree species distributions? Global Change Biology 24, 1150–1163.

25. Fait A, Angelovici R, Less H, Ohad I, Urbanczyk-Wochniak E, Fernie AR, Galili G. 2006. Arabidopsis Seed Development and Germination Is Associated with Temporally Distinct Metabolic Switches. Plant Physiology 142, 839–854.

26. Fang Z, Li M, Li J, et al. 2025. Identification of Malus sieversii ABA receptor PYL8 interacting proteome using Y2H-seq. Forestry Research 5.

27. Ferjani A, Segami S, Horiguchi G, Sakata A, Maeshima M, Tsukaya H. 2012. Regulation of pyrophosphate levels by H+-PPase is central for proper resumption of early plant development. Plant Signaling & Behavior 7, 38–42.

28. Fujii H, Verslues PE, Zhu J-K. 2011. Arabidopsis decuple mutant reveals the importance of SnRK2 kinases in osmotic stress responses in vivo. Proceedings of the National Academy of Sciences 108, 1717–1722.

29. Gallardo K, Job C, Groot S, Puype M, Demol H, Vandekerckhove J, Job D. 2002. Importance of methionine biosynthesis for Arabidopsis seed germination and seedling growth. Physiologia Plantarum 116, 238–247.

30. Ghanizadeh H, Qamer Z, Zhang Y, Wang A. 2025. The multifaceted roles of PP2C phosphatases in plant growth, signaling, and responses to abiotic and biotic stresses. Plant Communications 6, 101457.

31. Gipson AB, Morton KJ, Rhee RJ, et al. 2017. Disruptions in valine degradation affect seed development and germination in Arabidopsis. The Plant Journal 90, 1029–1039.

32. Gobert A, Isayenkov S, Voelker C, Czempinski K, Maathuis FJM. 2007. The two-pore channel TPK1 gene encodes the vacuolar K+ conductance and plays a role in K+ homeostasis. Proceedings of the National Academy of Sciences 104, 10726–10731.

33. Han G, Lu C, Guo J, Qiao Z, Sui N, Qiu N, Wang B. 2020. C2H2 Zinc Finger Proteins: Master Regulators of Abiotic Stress Responses in Plants. Frontiers in Plant Science 11.

34. Hang R, Xu Y, Wang X, Hu H, Flynn N, You C, Chen X. 2023. Arabidopsis HOT3/eIF5B1 constrains rRNA RNAi by facilitating 18S rRNA maturation. Proceedings of the National Academy of Sciences of the United States of America 120, e2301081120.

35. Harper AD, Bar-Peled M. 2002. Biosynthesis of UDP-Xylose. Cloning and Characterization of a Novel Arabidopsis Gene Family, UXS, Encoding Soluble and Putative Membrane-Bound UDP-Glucuronic Acid Decarboxylase Isoforms. Plant Physiology 130, 2188–2198.

36. Hasan MdM, Liu X-D, Waseem M, Guang-Qian Y, Alabdallah NM, Jahan MS, Fang X-W. 2022. ABA activated SnRK2 kinases: an emerging role in plant growth and physiology. Plant Signaling & Behavior 17, 2071024.

37. He Y, Galant A, Pang Q, Strul JM, Balogun SF, Jez JM, Chen S. 2011. Structural and Functional Evolution of Isopropylmalate Dehydrogenases in the Leucine and Glucosinolate Pathways of Arabidopsis thaliana. The Journal of Biological Chemistry 286, 28794–28801.

38. Hendrich R. 2012. Ion Channels in Plants | Physiological Reviews | American Physiological Society. Physiological Reviews 92, 1777–1811.

39. Hilhorst HW. 2007. Definitions and hypotheses of seed dormancy. In: Bradford KJ, Nonogaki H, eds. Seed Development, Dormancy and Germination. Oxford, UK: Blackwell, 50–71.

40. Hooks MA, Allwood JW, Harrison JKD, Kopka J, Erban A, Goodacre R, Balk J. 2014. Selective induction and subcellular distribution of ACONITASE 3 reveal the importance of cytosolic citrate metabolism during lipid mobilization in Arabidopsis. Biochemical Journal 463, 309–317.

41. Hu J. 2007. Toward Understanding Plant Peroxisome Proliferation. Plant Signaling & Behavior 2, 308–310.

42. Hu J, Baker A, Bartel B, Linka N, Mullen RT, Reumann S, Zolman BK. 2012. Plant Peroxisomes: Biogenesis and Function. The Plant Cell 24, 2279–2303.

43. Imhof J, Huber F, Reichelt M, Gershenzon J, Wiegreffe C, Lächler K, Binder S. 2014. The Small Subunit 1 of the Arabidopsis Isopropylmalate Isomerase Is Required for Normal Growth and Development and the Early Stages of Glucosinolate Formation. PLOS ONE 9, e91071.

44. ISTA. 2019. International Rules for Seed Testing. https://www.seedtest.org/en/publications/international-rules-seed-testing.html. Accessed January 2024.

45. Jagodzik P, Tajdel-Zielinska M, Ciesla A, Marczak M, Ludwikow A. 2018. Mitogen-Activated Protein Kinase Cascades in Plant Hormone Signaling. Frontiers in Plant Science 9.

46. Jiang Y, Wu X, Shi M, Yu J, Guo C. 2022. The miR159-MYB33-ABI5 module regulates seed germination in Arabidopsis. Physiologia Plantarum 174, e13659.

47. Jiménez JA, Rodríguez D, Lorenzo O, Nicolás G, Nicolás C. 2006. Characterization of a protein kinase (FsPK4) with an acidic domain, regulated by abscisic acid and specifically located in Fagus sylvatica L. seeds. Journal of Plant Physiology 163, 761–769.

48. Jin JB, Kim YA, Kim SJ, Lee SH, Kim DH, Cheong G-W, Hwang I. 2001. A New Dynamin-Like Protein, ADL6, Is Involved in Trafficking from the trans-Golgi Network to the Central Vacuole in Arabidopsis. The Plant Cell 13, 1511–1526.

49. Joseph MP, Papdi C, Kozma-Bognár L, Nagy I, López-Carbonell M, Rigó G, Koncz C, Szabados L. 2014. The Arabidopsis ZINC FINGER PROTEIN3 Interferes with Abscisic Acid and Light Signaling in Seed Germination and Plant Development1[C][W][OPEN]. Plant Physiology 165, 1203–1220.

50. Kanai M, Hayashi M, Kondo M, Nishimura M. 2013. The Plastidic DEAD-box RNA Helicase 22, HS3, is Essential for Plastid Functions Both in Seed Development and in Seedling Growth. Plant and Cell Physiology 54, 1431–1440.

51. Kao Y-T, Gonzalez KL, Bartel B. 2018. Peroxisome Function, Biogenesis, and Dynamics in Plants1[OPEN]. Plant Physiology 176, 162–177.

52. Kim JY. 2013. Identification and Functional Analysis of S-AdenosylMethionine Synthetase (HvSAMS) genes in Early Maturing Barley (Hordeum vulgare subsp. vulgare). Plant Breeding and Biotechnology 1, 178–195.

53. Kim P, Mahboob S, Nguyen HT, et al. 2024. Characterization of Soybean Events with Enhanced Expression of the Microtubule-Associated Protein 65-1 (MAP65-1). Molecular Plant-Microbe Interactions® 37, 62–71.

54. Klupczyńska EA, Pawłowski TA. 2021. Regulation of Seed Dormancy and Germination Mechanisms in a Changing Environment. International Journal of Molecular Sciences 22, 1357.

55. Kurepa J, Shull TE, Smalle JA. 2023. Friends in Arms: Flavonoids and the Auxin/Cytokinin Balance in Terrestrialization. Plants 12, 517.

56. Kurpisz B, Pawłowski TA. 2022. Epigenetic Mechanisms of Tree Responses to Climatic Changes. International Journal of Molecular Sciences 23, 13412.

57. Lächler K, Clauss K, Imhof J, Crocoll C, Schulz A, Halkier BA, Binder S. 2020. In Arabidopsis thaliana Substrate Recognition and Tissue- as Well as Plastid Type-Specific Expression Define the Roles of Distinct Small Subunits of Isopropylmalate Isomerase. Frontiers in Plant Science 11.

58. Langella O, Renne T, Balliau T, Davanture M, Brehmer S, Zivy M, Blein-Nicolas M, Rusconi F. 2024. Full Native timsTOF PASEF-Enabled Quantitative Proteomics with the i2MassChroQ Software Package. Journal of Proteome Research 23, 3353–3366.

59. Lee MB, Kim JY. 2024. HvVDAC1 interacts with HvSAMS1 and is predominantly expressed during germination and grain development. Journal of Plant Biochemistry and Biotechnology 33, 189–204.

60. Lefoulon C. 2021. The bare necessities of plant K+ channel regulation. Plant Physiology 187, 2092–2109.

61. Leuschner C, Ellenberg H. 2017. Ecology of Central European Forests: Vegetation Ecology of Central Europe, Volume I. Cham: Springer International Publishing.

62. Li X, Li C, Zhu J, Zhong S, Zhu H, Zhang X. 2023. Functions and mechanisms of RNA helicases in plants. Journal of Experimental Botany 74, 2295–2310.

63. Litterer LA, Plaisance KL, Schnurr JA, Storey KK, Jung H-JG, Gronwald JW, Somers DA. 2006. Biosynthesis of UDP-glucuronic acid in developing soybean embryos: possible role of UDP-sugar pyrophosphorylase. Physiologia Plantarum 128, 200–211.

64. Liu P-P, Montgomery TA, Fahlgren N, Kasschau KD, Nonogaki H, Carrington JC. 2007. Repression of AUXIN RESPONSE FACTOR10 by microRNA160 is critical for seed germination and post-germination stages. The Plant Journal 52, 133–146.

65. Liu L, Wang Y, Wang N, Dong Y-Y, Fan X-D, Liu X-M, Yang J, Li H-Y. 2011. Cloning of a Vacuolar H+-pyrophosphatase Gene from the Halophyte Suaeda corniculata whose Heterologous Overexpression Improves Salt, Saline-alkali and Drought Tolerance in Arabidopsis. Journal of Integrative Plant Biology 53, 731–742.

66. Lorenzo O, Nicolas C, Nicolas G, Rodriguez D. 2003. Characterization of a dual plant protein kinase (FsPK1) up-regulated by abscisic acid and calcium and specifically expressed in dormant seeds of Fagus sylvatica L. Seed Science Research 13, 261–271.

67. Lugassi N, Stein O, Egbaria A, Belausov E, Zemach H, Arad T, Granot D, Carmi N. 2022. Sucrose Synthase and Fructokinase Are Required for Proper Meristematic and Vascular Development. Plants 11, 1035.

68. Lyu T, Cao J. 2018. Cys2/His2 Zinc-Finger Proteins in Transcriptional Regulation of Flower Development. International Journal of Molecular Sciences 19, 2589.

69. Marchi M, Castellanos-Acuña D, Hamann A, Wang T, Ray D, Menzel A. 2020. ClimateEU, scale-free climate normals, historical time series, and future projections for Europe. Scientific Data 7, 428.

70. Markulin L, Corbin C, Renouard S, Drouet S, Gutierrez L, Mateljak I, Auguin D, Hano C, Fuss E, Lainé E. 2019a. Pinoresinol–lariciresinol reductases, key to the lignan synthesis in plants. Planta 249, 1695–1714.

71. Markulin L, Drouet S, Corbin C, et al. 2019b. The control exerted by ABA on lignan biosynthesis in flax (Linum usitatissimum L.) is modulated by a Ca2+ signal transduction involving the calmodulin-like LuCML15b. Journal of Plant Physiology 236, 74–87.

72. Meyer K, Stecca KL, Ewell-Hicks K, Allen SM, Everard JD. 2012. Oil and Protein Accumulation in Developing Seeds Is Influenced by the Expression of a Cytosolic Pyrophosphatase in Arabidopsis. Plant Physiology 159, 1221–1234.

73. Mravec J, Petrášek J, Li N, et al. 2011. Cell Plate Restricted Association of DRP1A and PIN Proteins Is Required for Cell Polarity Establishment in Arabidopsis. Current Biology 21, 1055–1060.

74. Muhammad D, Smith KA, Bartel B. 2022. Plant peroxisome proteostasis—establishing, renovating, and dismantling the peroxisomal proteome. Essays in Biochemistry 66, 229–242.

75. Nakashima K, Fujita Y, Kanamori N, et al. 2009. Three Arabidopsis SnRK2 Protein Kinases, SRK2D/SnRK2.2, SRK2E/SnRK2.6/OST1 and SRK2I/SnRK2.3, Involved in ABA Signaling are Essential for the Control of Seed Development and Dormancy. Plant and Cell Physiology 50, 1345–1363.

76. Nakatsubo T, Mizutani M, Suzuki S, Hattori T, Umezawa T. 2008. Characterization of Arabidopsis thaliana Pinoresinol Reductase, a New Type of Enzyme Involved in Lignan Biosynthesis*. Journal of Biological Chemistry 283, 15550–15557.

77. Nawaz G, Kang H. 2019. Rice OsRH58, a chloroplast DEAD-box RNA helicase, improves salt or drought stress tolerance in Arabidopsis by affecting chloroplast translation. BMC Plant Biology 19, 17.

78. Ogawa S, Mitsuya S. 2012. S-methylmethionine is involved in the salinity tolerance of Arabidopsis thaliana plants at germination and early growth stages. Physiologia Plantarum 144, 13–19.

79. Pan C, Yao L, Yu L, Qiao Z, Tang M, Wei F, Huang X, Zhou Y. 2023. Transcriptome and proteome analyses reveal the potential mechanism of seed dormancy release in Amomum tsaoko during warm stratification. BMC Genomics 24, 99.

80. Pang X, Liu S, Suo J, et al. 2023. Proteome Dynamics Analysis Reveals the Potential Mechanisms of Salinity and Drought Response during Seed Germination and Seedling Growth in Tamarix hispida. Genes 14.

81. Pawłowski TA. 2009. Proteome analysis of Norway maple (Acer platanoides L.) seeds dormancy breaking and germination: influence of abscisic and gibberellic acids. BMC Plant Biology 9, 48.

82. Pawłowski TA. 2010. Proteomic approach to analyze dormancy breaking of tree seeds. Plant Molecular Biology 73, 15–25.

83. Pawłowski TA, Bergervoet JHW, Bino RJ, Groot SPC. 2004. Cell cycle activity and β-tubulin accumulation during dormancy breaking of Acer platanoides L. seeds. Biologia Plantarum 48, 211–218.

84. Pawłowski TA, Bujarska-Borkowska B, Suszka J, Tylkowski T, Chmielarz P, Klupczyńska EA, Staszak AM. 2020. Temperature regulation of primary and secondary seed dormancy in Rosa canina L.: findings from proteomic analysis. International Journal of Molecular Sciences 21, 7008.

85. Pawłowski TA, Staszak AM. 2016. Analysis of the embryo proteome of sycamore (Acer pseudoplatanus L.) seeds reveals a distinct class of proteins regulating dormancy release. Journal of Plant Physiology 195, 9–22.

86. Pawłowski TA, Suszka J. 2025. Proteome analysis provides insight into dormancy and germination of silver fir embryo and megagametophyte. Dendrobiology 94, 46–61.

87. Pawłowski TA, Suszka J, Mucha J, Zadworny M, Alipour S, Kurpisz B, Chmielarz P, Jagodziński AM, Chmura DJ. 2024. Climate legacy in seed and seedling traits of European beech populations. Frontiers in Plant Science 15.

88. Penfield S. 2017. Seed dormancy and germination. Current Biology 27, R874–R878.

89. R Core Team. 2024. R: A Language and Environment for Statistical Computing.

90. Rahikainen M, Berkowitz O, Whelan J, Kangasjärvi S, Pascual J. 2025. Role of aconitase in plant stress response and signaling. Physiologia Plantarum 177, e70128.

91. Ramachandran V, Chen X. 2008. Degradation of microRNAs by a Family of Exoribonucleases in Arabidopsis. Science 321, 1490–1492.

92. Reboul R, Geserick C, Pabst M, Frey B, Wittmann D, Lütz-Meindl U, Léonard R, Tenhaken R. 2011. Down-regulation of UDP-glucuronic Acid Biosynthesis Leads to Swollen Plant Cell Walls and Severe Developmental Defects Associated with Changes in Pectic Polysaccharides*. Journal of Biological Chemistry 286, 39982–39992.

93. Ren S, Lv G. 2024. A Proteomic Study on Seed Germination of Nitraria roborowskii Kom. Forests 15, 1661.

94. Renouard S, Corbin C, Lopez T, Montguillon J, Gutierrez L, Lamblin F, Lainé E, Hano C. 2012. Abscisic acid regulates pinoresinol–lariciresinol reductase gene expression and secoisolariciresinol accumulation in developing flax (Linum usitatissimum L.) seeds. Planta 235, 85–98.

95. Reyes JL, Chua N-H. 2007. ABA induction of miR159 controls transcript levels of two MYB factors during Arabidopsis seed germination. The Plant Journal 49, 592–606.

96. Reyes D, Rodríguez D, Lorenzo O, Nicolás G, Cañas R, Cantón FR, Canovas FM, Nicolás C. 2006. Immunolocalization of FsPK1 correlates this abscisic acid-induced protein kinase with germination arrest in Fagus sylvatica L. seeds. Journal of Experimental Botany 57, 923–929.

97. Rinne J, Niehaus M, Medina-Escobar N, Straube H, Schaarschmidt F, Rugen N, Braun H-P, Herde M, Witte C-P. 2024. Three Arabidopsis UMP kinases have different roles in pyrimidine nucleotide biosynthesis and (deoxy)CMP salvage. The Plant Cell 36, 3611–3630.

98. Roje S. 2006. S-Adenosyl-l-methionine: Beyond the universal methyl group donor. Phytochemistry 67, 1686–1698.

99. Rosbakh S, Pacini E, Nepi M, Poschlod P. 2018. An Unexplored Side of Regeneration Niche: Seed Quantity and Quality Are Determined by the Effect of Temperature on Pollen Performance. Frontiers in Plant Science 9.

100. Sajeev N, Koornneef M, Bentsink L. 2024. A commitment for life: Decades of unraveling the molecular mechanisms behind seed dormancy and germination. The Plant Cell 36, 1358–1376.

101. Sano N, Lounifi I, Cueff G, Collet B, Clément G, Balzergue S, Huguet S, Valot B, Galland M, Rajjou L. 2022. Multi-Omics Approaches Unravel Specific Features of Embryo and Endosperm in Rice Seed Germination. Frontiers in Plant Science 13.

102. Sauter M, Moffatt B, Saechao MC, Hell R, Wirtz M. 2013. Methionine salvage and S-adenosylmethionine: essential links between sulfur, ethylene and polyamine biosynthesis. Biochemical Journal 451, 145–154.

103. Sawa S, Koizumi K, Naramoto S, Demura T, Ueda T, Nakano A, Fukuda H. 2005. DRP1A Is Responsible for Vascular Continuity Synergistically Working with VAN3 in Arabidopsis. Plant Physiology 138, 819–826.

104. Schachtman DP. 2000. Molecular insights into the structure and function of plant K+ transport mechanisms. Biochimica et Biophysica Acta (BBA) - Biomembranes 1465, 127–139.

105. Segami S, Asaoka M, Kinoshita S, Fukuda M, Nakanishi Y, Maeshima M. 2018. Biochemical, Structural and Physiological Characteristics of Vacuolar H+-Pyrophosphatase. Plant and Cell Physiology 59, 1300–1308.

106. Sghaier-Hammami B, B. M. Hammami S, Baazaoui N, Gómez-Díaz C, Jorrín-Novo JV. 2020. Dissecting the Seed Maturation and Germination Processes in the Non-Orthodox Quercus ilex Species Based on Protein Signatures as Revealed by 2-DE Coupled to MALDI-TOF/TOF Proteomics Strategy. International Journal of Molecular Sciences 21, 4870.

107. Sharma T, Dreyer I, Riedelsberger J. 2013. The role of K+ channels in uptake and redistribution of potassium in the model plant Arabidopsis thaliana. Frontiers in Plant Science 4.

108. Shirley BW, Hanley S, Goodman HM. 1992. Effects of ionizing radiation on a plant genome: analysis of two Arabidopsis transparent testa mutations. The Plant Cell 4, 333–347.

109. Singh R, Iqbal N, Umar S, Ahmad S. 2024. Lignan Enhancement: An Updated Review on the Significance of Lignan and Its Improved Production in Crop Plants. Phyton-International Journal of Experimental Botany 93, 3237–3271.

110. Smertenko AP, Chang H-Y, Sonobe S, Fenyk SI, Weingartner M, Bögre L, Hussey PJ. 2006. Control of the AtMAP65-1 interaction with microtubules through the cell cycle. Journal of Cell Science 119, 3227–3237.

111. Stein O, Granot D. 2018. Plant Fructokinases: Evolutionary, Developmental, and Metabolic Aspects in Sink Tissues. Frontiers in Plant Science 9, 339.

112. Stephen JR, Dent KC, Finch-Savage WE. 2004. Molecular responses of Prunus avium (wild cherry) embryonic axes to temperatures affecting dormancy. New Phytologist 161, 401–413.

113. Tan H, Man C, Xie Y, Yan J, Chu J, Huang J. 2019. A Crucial Role of GA-Regulated Flavonol Biosynthesis in Root Growth of Arabidopsis. Molecular Plant 12, 521–537.

114. Tang H, Vasconcelos AC, Berkowitz GA. 1996. Physical association of KAB1 with plant K+ channel alpha subunits. The Plant Cell 8, 1545–1553.

115. Taylor NL, Heazlewood JL, Day DA, Millar AH. 2004. Lipoic Acid-Dependent Oxidative Catabolism of α-Keto Acids in Mitochondria Provides Evidence for Branched-Chain Amino Acid Catabolism in Arabidopsis. Plant Physiology 134, 838–848.

116. Tognacca RS, Botto JF. 2021. Post-transcriptional regulation of seed dormancy and germination: Current understanding and future directions. Plant Communications 2, 100169.

117. Valot B, Langella O, Nano E, Zivy M. 2011. MassChroQ: a versatile tool for mass spectrometry quantification. Proteomics 11, 3572–3577.

118. Vavrdová T, _̌_Samaj J, Komis G. 2019. Phosphorylation of Plant Microtubule-Associated Proteins During Cell Division. Frontiers in Plant Science 10, 238.

119. Wang X, Davanture M, Zivy M, et al. 2022a. Label-Free Quantitative Proteomics Reveal the Involvement of PRT6 in Arabidopsis thaliana Seed Responsiveness to Ethylene. International Journal of Molecular Sciences 23.

120. Wang X, Davanture M, Zivy M, Bailly C, Nambara E, Corbineau F. 2022b. Label-Free Quantitative Proteomics Reveal the Involvement of PRT6 in Arabidopsis thaliana Seed Responsiveness to Ethylene. International Journal of Molecular Sciences 23, 9352.

121. Wang J, Johnson AG, Lapointe CP, Choi J, Prabhakar A, Chen D-H, Petrov AN, Puglisi JD. 2019. eIF5B gates the transition from translation initiation to elongation. Nature 573, 605–608.

122. Wang Y, Liao Y, Quan C, et al. 2022c. C2H2-type zinc finger OsZFP15 accelerates seed germination and confers salinity and drought tolerance of rice seedling through ABA catabolism. Environmental and experimental botany in press.

123. Wang H, Wang J, Guo X, Brennan CS, Li T, Fu X, Chen G, Liu RH. 2016a. Effect of germination on lignan biosynthesis, and antioxidant and antiproliferative activities in flaxseed (Linum usitatissimum L.). Food Chemistry 205, 170–177.

124. Wang Y-M, Yang Q, Liu Y-J, Yang H-L. 2016b. Molecular Evolution and Expression Divergence of the Aconitase (ACO) Gene Family in Land Plants. Frontiers in Plant Science 7, 1879.

125. Weijers D, Franke-van Dijk M, Vencken R-J, Quint A, Hooykaas P, Offringa R. 2001. An Arabidopsis Minute-like phenotype caused by a semi-dominant mutation in a RIBOSOMAL PROTEIN S5 gene. Development 128, 4289–4299.

126. Weis BL, Missbach S, Marzi J, Bohnsack MT, Schleiff E. 2014. The 60S associated ribosome biogenesis factor LSG1-2 is required for 40S maturation in Arabidopsis thaliana. The Plant Journal 80, 1043–1056.

127. Wickham H. 2016. ggplot2: Elegant Graphics for Data Analysis. New York: Springer-Verlag.

128. Wickham H. 2023. stringr: Simple, Consistent Wrappers for Common String Operations. R package version 1.5.1.

129. Wickham H, Francois R., Henry L., Muller K., Vaughan D. 2023. dplyr: A Grammar of Data Manipulation. R package version 1.3.1.

130. Wickham H, Vaughan D, Girlich M. 2024. tidyr: Tidy Messy Data. R package version 1.3.1.

131. Wingenter K, Trentmann O, Winschuh I, Hörmiller II, Heyer AG, Reinders J, Schulz A, Geiger D, Hedrich R, Neuhaus HE. 2011. A member of the mitogen-activated protein 3-kinase family is involved in the regulation of plant vacuolar glucose uptake. The Plant Journal 68, 890–900.

132. Wojtyla Ł, Lechowska K, Kubala S, Garnczarska M. 2016. Different Modes of Hydrogen Peroxide Action During Seed Germination. Frontiers in Plant Science 7.

133. Wu B, Liang H, Lv J, Liu R, Ye N. 2025. Key Regulators of Seed Germination: Kinases and Phosphatases. Seeds 4, 30.

134. Xia Q, Maharajah P, Cueff G, Rajjou L, Prodhomme D, Gibon Y, Bailly C, Corbineau F, Meimoun P, El-Maarouf-Bouteau H. 2018. Integrating proteomics and enzymatic profiling to decipher seed metabolism affected by temperature in seed dormancy and germination. Plant Science 269, 118–125.

135. Yan H, Chaumont N, Gilles JF, Bolte S, Hamant O, Bailly C. 2020. Microtubule self-organisation during seed germination in Arabidopsis. BMC Biology 18, 44.

136. Yoshida T, Nishimura N, Kitahata N, Kuromori T, Ito T, Asami T, Shinozaki K, Hirayama T. 2006. ABA-hypersensitive germination3 encodes a protein phosphatase 2C (AtPP2CA) that strongly regulates abscisic acid signaling during germination among Arabidopsis protein phosphatase 2Cs. Plant physiology 140, 115–126.

137. Yu Y, Jia T, Chen X. 2017. The ‘how’ and ‘where’ of plant microRNAs. New Phytologist 216, 1002–1017.

138. Yuan P, Cai Q, Hu Z. 2024. Arabidopsis DEAD-box RNA helicase 12 is required for salt tolerance during seed germination. Biochemical and Biophysical Research Communications 725, 150228.

139. Zhang L, Liu X, Gaikwad K, et al. 2017. Mutations in eIF5B Confer Thermosensitive and Pleiotropic Phenotypes via Translation Defects in Arabidopsis thaliana. The Plant Cell 29, 1952–1969.

140. Zhang Q, Shirley N, Lahnstein J, Fincher GB. 2005. Characterization and Expression Patterns of UDP-d-Glucuronate Decarboxylase Genes in Barley. Plant Physiology 138, 131–141.

141. Zhang M, Zhang Y, Deng Z, Liu T, Liang Y, Li Q, Zou C, Chen Z, Ma L, Shen Y. 2025. GWAS and gene co-expression network analysis reveal the genetic control of seed germination under salt stress in maize. Theoretical and Applied Genetics 138, 285.

142. Zhou L, Lacroute F, Thornburg R. 1998. Cloning, Expression in Escherichia coli, and Characterization of Arabidopsis thaliana UMP/CMP Kinase1. Plant Physiology 117, 245–254.

